# Conserved Metastatic Cell States and Spatial Immune Microenvironment Remodeling Define Pancreatic Cancer Liver Metastases

**DOI:** 10.64898/2026.05.27.728281

**Authors:** Ayushi Mandloi, Christina R. Larson, Tuan M. Tran, Sudharonjon Roy, Ateeq M. Khaliq, Yu-Hua Dean Fang, Jace Baines, Richard P. Laube, Meet Patel, Christopher Risley, Robert S. Welner, Satwik Acharyya, Ashiq Masood, Julienne L. Carstens

## Abstract

**Background:** Pancreatic ductal adenocarcinoma (PDAC) mortality is driven largely by liver metastatic disease; however, the malignant cell states and tumor microenvironmental features of PDAC liver metastases remain obscure.

**Methods:** We developed a transplant model system of matched pancreatic and liver tumors to study PDAC metastatic progression. Using this model, we identified murine PDAC cell lines with distinct liver metastatic capacities and performed multiomic profiling of matched primary pancreatic and metastatic liver tumors. Transcriptional programs associated with high and low liver tropism were defined and evaluated across tumor models and independent human PDAC datasets. Spatial and tumor-immune interaction analyses were used to characterize microenvironmental niches, immune composition, and cellular relationships within primary and metastatic tumors.

**Results:** A high-liver-tropic transcriptional program was enriched in high liver-tropic cell lines and malignant cells within liver metastases, conserved across human PDAC datasets, and associated with inferior patient survival. High- and low-liver-tropic tumor states occupied distinct liver microenvironmental niches and exhibited different tumor-immune communication networks, accompanied by local and systemic changes in immune composition. Liver metastases also displayed features of enhanced immunosuppression, including increased proximity of CD4^+^ and CD8^+^ T cells to tumor cells and greater spatial association between regulatory T cells and exhausted CD8^+^ T cells.

**Conclusions:** These findings identify conserved PDAC cell states associated with differential liver metastatic capacity and demonstrate that liver metastasis is accompanied by spatial and immunologic remodeling of the tumor microenvironment. Together, the study provides a framework for understanding how tumor-intrinsic metastatic programs interact with site-specific immune ecosystems in PDAC.

## Background

Metastatic PDAC is the dominant clinical presentation, with only a 3% 5-year survival rate (1). Over 80% of metastatic disease occurs in the liver and contributes substantially to this poor prognosis (2–5). Recent clinical studies have demonstrated that primary tumors harbor distinct transcriptional programs associated with liver or lung metastatic tropism (6). The primary tumors with enhanced liver tropism exhibited elevated tolerance to replication stress, while the tumors with reduced liver tropism were associated with increased lymphocyte densities and T cell clonal responses (6). These findings suggest that cancer-intrinsic programs and interactions within the primary tumor microenvironment (TME) influence metastatic organotropism. However, the role of the metastatic site microenvironment in determining this organotropism remains unexplored.

Recent studies have revealed that metastatic lesions develop within tissue-specific TME that differ substantially from their primary tumor counterparts. Spatial transcriptomic analyses of matched primary and metastatic PDAC specimens have demonstrated distinct tumor ecosystems across anatomical sites (7,8). Notably, we identified a complex mixture of pro- and suppressive-immune signatures at the invasive edge of liver metastases, suggesting that a unique tumor-immune interaction may facilitate metastatic outgrowth in the liver (7). Other studies, largely using unmatched samples, have also highlighted the differences in immune composition and organization between pancreatic and liver metastatic TMEs (9). Together, these observations support the existence of site-specific immune ecosystems in PDAC metastasis but remain largely descriptive, limiting our ability to determine how malignant cell programs and local immune environments interact during metastatic progression.

The Kras- and p53-mutant-driven (KPC) genetically engineered mouse models (GEMM) represent nearly 60% of clinical PDAC cases and are a foundational pre-clinical platform for studying pancreatic cancer progression. Studies using this model have revealed that liver metastatic progression is associated with distinct TME features, including increased desmoplasia with lesion growth, while spontaneous liver metastases have been linked to epithelial characteristics of the primary tumor (10–13). However, reports using KPC GEMMs have varied substantially in the frequency and extent of metastatic disease, with some studies describing predominantly microscopic and small metastatic lesions, while others report more frequent overt metastatic burden (11,14). Because patients with metastatic PDAC are diagnosed based on clinically detectable lesions through clinical imaging (approximately 10 mm) rather than microscopic disease (< 2 mm), experimental systems capable of generating robust and reproducible metastatic outgrowth may better facilitate interrogation of clinically relevant metastatic biology (15).

To circumvent the KPC model’s rate-limiting step of primary tumor intravasation, liver colonization models using direct or indirect portal vein delivery have become increasingly common (16,17). Early studies using hemisplenic injection demonstrated efficient hepatic outgrowth but did not interrogate the cancer cell-intrinsic or microenvironmental factors associated with this process (18). Furthermore, the use of different cell lines with variable engraftment rates has made it difficult to distinguish intrinsic liver metastatic capacity from site-specific microenvironmental influences. Finally, these approaches lack a matched primary tumor, limiting direct comparisons between primary and metastatic disease and reducing their relevance to stage IV clinical presentation.

In this study, we established a dual transplant system that generates matched pancreatic and liver tumors using primary tumor-derived syngeneic KPC cell lines in immunocompetent mice. Leveraging this platform, we identified distinct metastatic phenotypes associated with differential liver outgrowth and defined a transcriptional signature enriched in metastatic tumors. This metastatic signature was conserved across multiple human PDAC datasets and was enriched in malignant cells from liver metastases. Integrative analyses further revealed site-specific tumor microenvironments characterized by altered immune signaling networks, distinct spatial organization of immune cells relative to cancer cells, and systemic remodeling of immune cells with progressive metastasis. Collectively, these studies identify conserved metastatic cell states and spatial immune remodeling associated with PDAC liver metastases while providing an experimentally tractable framework for investigating mechanisms that govern metastatic progression and therapeutic response.

## Results

### PDAC cell-intrinsic programs associate with differential liver metastatic outgrowth

The organ-specific metastatic patterns observed in PDAC are thought to arise from complex interactions between cancer cell-intrinsic programs and the permissive properties of the distant tissue microenvironments (19). To investigate whether cancer cell-intrinsic properties contribute to the differential liver metastatic potential within this model system, we characterized six C57Bl/6 syngeneic KPC primary tumor-derived cell lines spanning the epithelial-to-mesenchymal transition (EMT) spectrum (**Figure 1A** and **Supplementary Tables 1 and 2**).

**Figure 1:**
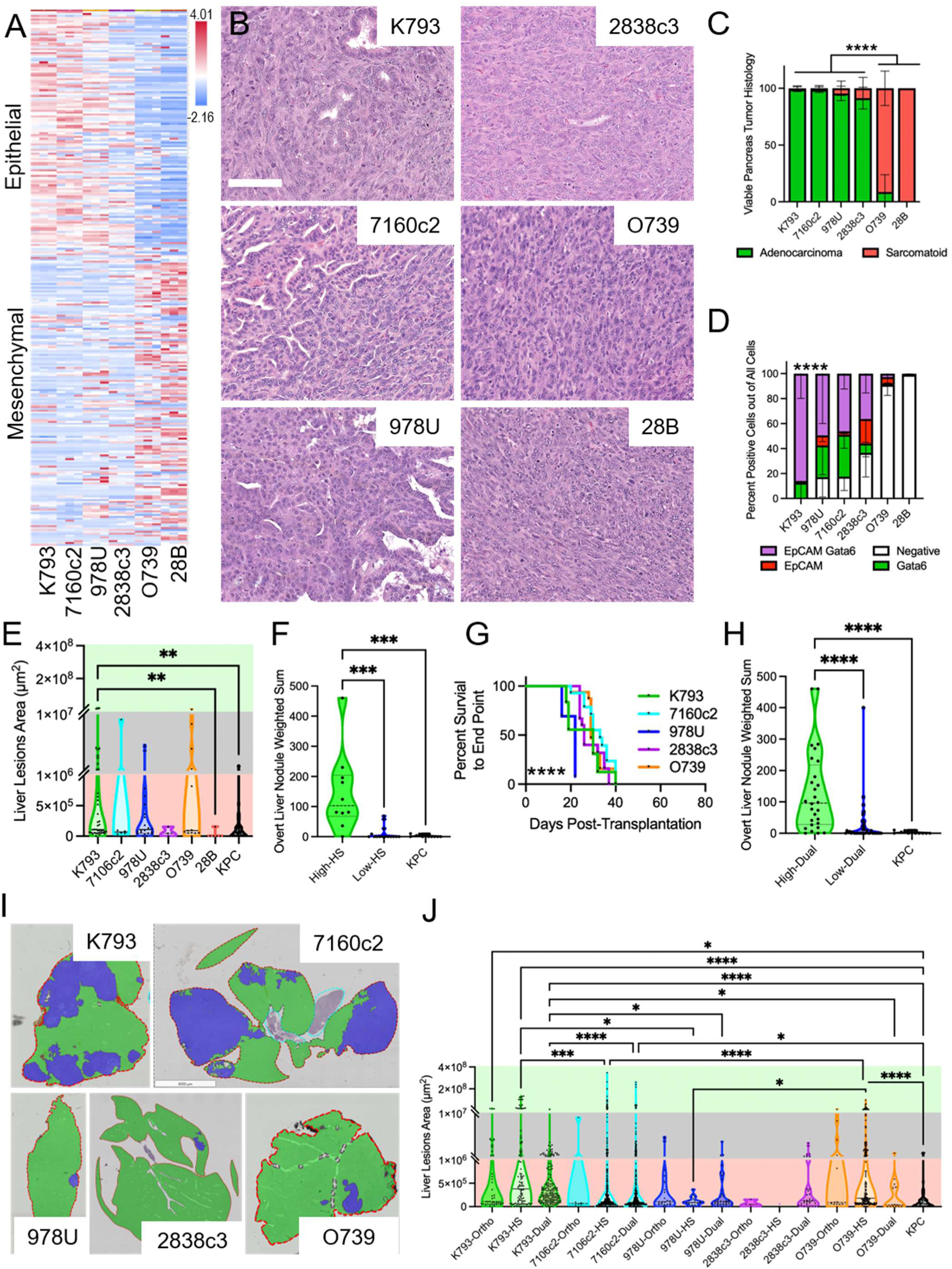
Cancer cell-intrinsic differences define liver metastatic capacity. **A**. RNA-seq Z-score of a 315 universal EMT signature (26) across the indicated cell lines. **B**. Representative histopathological micrographs of orthotopic transplants (scale 100 µm). **C**. Relative percentage of histological subtype within the viable tumor regions of orthotopic pancreatic tumors. n = K793:7, 7160c2: 9, 978U: 7, 2838c3: 6, O739: 6, 28B: 3 mice. Two-way ANOVA, multiple comparisons showed each of the predominantly adenocarcinoma lines was significantly different from each of the sarcomatoid lines; a summary comparison was graphed for simplicity of display. **D**. Quantification of epithelial (EpCam) and Classical (Gata6) markers in YFP^+^ cancer cells from orthotopic pancreatic tumors. n = 3 mice for each line. Two-way ANOVA. **E.** Distribution of liver metastatic lesion areas following orthotopic transplantation of the indicated cell lines. Axis breaks correspond to clinical imaging groups of micrometastases (red), borderline detectable (gray), and detectable (green). **F**. Weighted sum by a binned size of overt liver nodules of hemisplenic (HS) injected cells (100,000) compared to the spontaneous KPC mouse model. n = High: 10, Low: 15, KPC: 11. **G**. Percent survival to endpoint of dual transplants. n = K793: 17, 7160c2: 13, 978U: 12, 2838c3: 14, O739: 14. Mantel-Cox test. **H**. Weighted sum by binned size of overt liver nodules of dual-injected cells compared to the spontaneous KPC mouse model. n = High: 28, Low: 38, KPC: 11. **I**. Representative dual liver scans with tumor (blue) and normal liver (green) overlays (scale 6500 µm). **J**. Quantification of each metastatic liver nodule area of the indicated cell lines and injections. Axis breaks correspond to clinical imaging groups of micrometastases (red), borderline detectable (gray), and detectable (green). Significance determined by Kruskal-Wallis One-Way ANOVA, unless indicated otherwise. * p<0.05, ** p<0.01, *** p<0.001, **** p<0.0001, ns, not significant.

Orthotopic transplantation preserved the adenocarcinoma and sarcomatoid morphologies predicted by transcriptional EMT status and observed in spontaneous KPC tumors (**Figures 1B** and **1C** and **Supplementary Figures 1A and 1B**). Consistent with established relationships between EMT and classical/basal transcriptional programs, expression of Gata6 and EpCAM tracked with the morphological and EMT status of each line (**Figure 1D** and **Supplementary Figures 1C**) (20,21). Liver metastasis varied by cell line as the size and incidence of overt nodules was comparable to the KPC GEMMs; however, microscopic lesion area showed significant differences. (**Figure 1E**, **Supplementary Figure 1D,** and **Supplementary** Tables 3 and 4) (22). These findings demonstrated cell line-dependent differences in spontaneous liver metastatic burden, reflecting variability in both metastatic seeding and outgrowth.

To determine whether these differences reflected cancer cell-intrinsic metastatic capacity, we compared liver outgrowth across the lineage-traced KPC cell line panel using hemisplenic transplantation (23). Two cell lines, K793 and 7160c2, consistently generated overt liver tumors, whereas the remaining lines exhibited limited metastatic outgrowth (**Supplementary Table 3**). Pooling these high liver-tropic phenotype cell lines showed a significant increase in overt lesion size compared to the low liver-tropic phenotype cell lines and KPC GEMMs (**Figure 1F**). The definition of high and low liver tropism spans cell lines from differing genetic backgrounds and genotypes, supporting their metastatic potential as cancer-cell intrinsic and not an experimental artifact. Altering the number of injected cells had minimal effect on liver lesion size in the high-tropic K793 line, whereas low-tropic lines (978U and 2838c3) displayed only modest increases in lesion burden at the highest cell doses. Even under these conditions, lesions largely remained microscopic or borderline detectable, indicating that differential metastatic outgrowth was not solely explained by initial tumor cell number (24,25) (**Supplementary Figure 1E**). Direct portal vein injection yielded similar relative differences between high- and low-tropic lines but introduced experimental engraftment artifacts in the collagen and growth-factor-rich hemostatic gauze, supporting continued use of the hemisplenic approach (17) (**Supplementary Figure 1F**). Altogether, these data further support the use of the hemisplenic transplantation model for liver tumor seeding and outgrowth; however, its performance is heavily cell line dependent, showing high- and low-liver tropism, and still lacks a matched pancreas/primary tumor to replicate the clinical presentation of PDAC.

To enable direct comparison of a matched pancreas and liver tumors within the same host, we combined the orthotopic and hemisplenic transplantation in a single surgical session, referred to as the dual transplant system. Importantly, the high- and low-liver-tropic phenotypes observed in single-site transplantation were preserved, with liver tumor incidence and metastatic burden closely mirroring those observed in the hemisplenic model. Transplantation rates in the pancreas either improved or remained consistent according to cell line, revealing a potential synergy between the tumor sites (**Supplementary Table 3**). With the addition of liver tumor burden, the time to endpoint (defined by large tumor size, drop in body condition, or moribundity) was shortened for slower-growing lines compared to orthotopic transplants, thereby improving the consistency of this model for time-course studies (**Figure 1G** and **Supplementary Figure 1G**). The pancreatic tumors retained the cellular morphology and histopathology scores observed in single orthotopic transplants (**Supplementary Figure 1H**). Liver tumor incidence and lesion size were like those of the hemisplenic models and retained the high and low-liver tropism previously observed (**Figures 1H-J** and **Supplementary Tables 3 and 4**). Collectively, these studies identify distinct liver metastatic cell states across KPC-derived PDAC cell lines and demonstrate that these states are maintained in matched pancreatic and liver metastatic tumors, enabling direct investigation of site-specific features of metastatic progression.

### Cross-species analyses identify a conserved PDAC liver metastatic signature

Next, we sought to identify cancer cell-intrinsic transcriptional programs associated with these distinct metastatic cell states. Bulk RNA-sequencing of the KPC cell lines identified 462 genes commonly upregulated in the high liver-tropic lines (K793 and 7160c2) compared to the low liver-tropic lines (978U, 2838c3, and O739), and 47 genes were enriched in the low liver-tropic lines relative to their high-tropic counterparts (**Figure 2A**, **Supplementary Tables 5 and 6**). Functional enrichment analysis using Metascape revealed that genes associated with high liver tropism were predominantly enriched for metabolic and mitochondrial processes, matching other reports of how metabolism can influence metastatic tumor cell tropism and outgrowth (**Supplementary Figure 2A**) (27–32). Conversely, genes enriched in low liver-tropic lines were associated with cellular stress responses, inflammatory signaling, and regulation of cell adhesion (**Supplementary Figure 2B**). Collectively, these analyses identified a metastatic-associated transcriptional program characterized by enhanced metabolic and mitochondrial pathways and reduced inflammatory and stress-response signaling.

**Figure 2:**
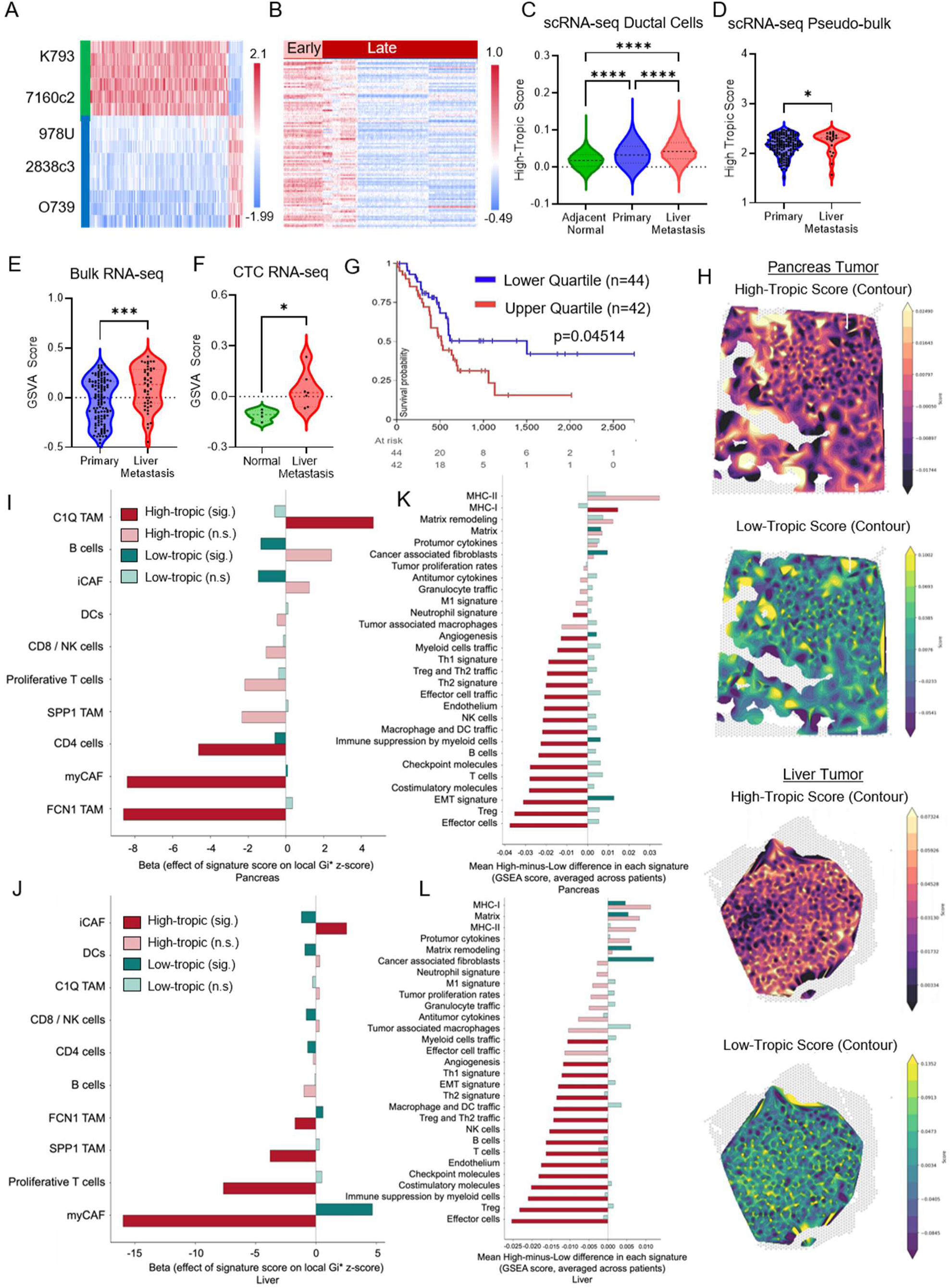
A Conserved Metastatic Cell State Is Enriched in PDAC Liver Metastases. **A.** RNA-seq Z-score of High- (green) and low-tropic (blue) gene signatures. **B**. scRNA-seq Z-score of High-Tropic signature in spontaneous KPC cancer cells in the early and late liver tumors. **C-D.** High liver-tropic score in all ductal cells (C) or pseudo-bulked of all single cells by sample (D) from a pooled patient-derived scRNA-seq data set. Kruskal-Wallis One-Way ANOVA with Dunn’s multiple comparisons or a Mann-Whitney T test, respectively. **E-F**. GSVA score of the High-Tropic signature in bulk RNA-seq of all samples from a rapid autopsy sample (E) and circulating epithelial cells from normal/healthy controls or patients with PDAC liver metastasis (F). Unpaired T test. **G**. Kaplan-Meier overall survival analysis of TCGA-PAAD patients stratified by upper and lower quartiles of expression of the High-liver-tropic gene signature. **H.** Smoothed contour maps of spatial transcriptomic sections from patient-derived primary pancreatic tumor and liver metastases indicating areas within the metastasis-associated ecotypes enriched for the high- and low-liver-tropic signature **I-J**. Cell-type composition of the metastasis-associated ecotypes according to high- and low-liver-tropic gene signature for patient primary pancreatic tumors (I) and liver metastases (J). **K-L.** Functional gene expression signatures (FGES) comparing high- and low-liver-tropic signature positive spots in metastasis-associated ecotypes from patient primary pancreatic tumors (K) and liver metastases (L). * p<0.05, *** p<0.001, **** p<0.0001, ns, not significant.

We next evaluated whether this high liver-tropic transcriptional program was associated with metastatic progression *in vivo*. In a previously published scRNA-seq dataset containing matched KPC primary tumors and liver metastases, the signature was enriched in a subset of metastatic cells, particularly among liver-disseminated populations from early, precancerous pancreatic lesions (**Figure 2B** and **Supplementary Figure 2C**) (12). We then examined a human PDAC scRNA-seq atlas and found significant enrichment of the signature in ductal cells from liver metastases compared with primary tumors and normal pancreata (**Figure 2C** and **Supplementary Figure 2D**) (33). These findings suggest that the signature is associated with early metastatic populations during liver colonization.

To determine whether this enrichment persisted at the patient level, we pseudo-bulked cells by patient and found that liver metastases retained higher signature expression than primary tumors (**Figure 2D**), consistent with increased enrichment in metastatic lesions by GSVA of rapid-autopsy bulk RNA-seq cohorts. (**Figure 2E**) (34). This pattern persisted when samples were averaged by patient and when analyses were restricted to matched primary-metastatic pairs (**Supplementary Figures 2E-F**). To assess disease specificity, we evaluated the high-liver-tropic signature across breast, colorectal, and hepatocellular carcinoma transcriptomic datasets, including liver metastases where available. In contrast to PDAC, the signature was not significantly enriched in these malignancies, suggesting that it represents a PDAC-specific liver metastatic program (**Supplementary Figure 2G-K**).

To determine whether this program could be detected in circulation, thereby helping detect occult recurrence, we analyzed circulating tumor cell (CTC) datasets and confirmed increased enrichment in patients with metastatic liver disease compared to healthy controls (**Figure 2F**) (35). We next assessed whether the expression of the high liver-tropic signature was associated with patient survival outcomes. Stratification of the primary tumor samples in the TCGA PAAD patients by upper and lower quartiles of signature expression demonstrated significantly reduced overall survival in patients with high signature expression (**Figure 2G**).

To investigate the spatial microenvironment associated with these distinct high and low metastatic cell states, we analyzed patient-derived spatial transcriptomic datasets and narrowed our assessment to the previously defined liver metastasis-associated ecotypes CC2 and CC3 (7). Smoothed contour mapping of high- and low-liver tropic signature scores revealed largely non-overlapping spatial territories, indicating that these programs occupy distinct microenvironmental niches (**Figure 2H**). Comparison of cellular associations within these regions demonstrated that in primary pancreatic tumors, myCAF signatures were significantly depleted in high-liver-tropic regions and enriched in low-liver-tropic regions, whereas CD4^+^ T-cell signatures were significantly reduced in both programs (**Figure 2I**). A similar pattern was observed in liver metastases, where myCAF signatures remained depleted in high-liver-tropic regions and enriched in low-liver-tropic regions. In addition, high-liver-tropic regions were enriched for iCAF signatures and depleted of FCN1 TAM signatures, while low-liver-tropic regions showed the opposite pattern (**Figure 2J**).

To determine whether these spatially distinct regions also differed functionally, we compared published functional gene expression signatures (FGES). Across both primary pancreatic tumors and liver metastases, high-liver tropic regions were characterized by reduced immune-related signatures, including T cells, B cells, NK cells, myeloid cells, and immune checkpoint pathways, whereas low-tropic regions were enriched for fibroblast and extracellular matrix-related programs (**Figures 2K–L**). Together, these findings indicate that high- and low-liver-tropic programs define conserved spatial niches across primary pancreatic tumors and liver metastases, distinguished by distinct stromal organization and reduced immune-associated features within the high-liver-tropic program.

Collectively, these findings identify a conserved metastatic cell state that is enriched across experimental and human PDAC liver metastases and associated with adverse clinical outcomes and microenvironmental ecosystems. These data support the hypothesis that this transcriptional program guides site-specific microenvironmental organization and metastatic progression.

### Conserved tumor–immune signaling programs characterize PDAC liver metastases

To determine whether these metastatic cell states have the potential to communicate with their tumor–microenvironment, we used NicheNet to predict ligand-receptor interactions in published human and spontaneous murine scRNA-seq datasets containing primary pancreatic tumors and liver metastases (12,33). Of the 462 genes in the high-tropic signature, 18 were annotated as NicheNet ligands, 11 of which were predicted to influence microenvironmental cells (**Supplementary Table 5**), suggesting a potential role for liver-tropic cancer cells in shaping the metastatic microenvironment.

In human liver metastases, predicted cancer cell-CD8^+^ T-cell interactions were ranked most highly for TGFB1-, LGALS3-, and SPP1-associated networks relative to primary tumors (**Figures 3A-3B**). These pathways have been implicated in immune suppression, myeloid-cell recruitment, and extracellular matrix remodeling in PDAC (36–42). Similar signaling programs were observed in GEMMs, alongside enrichment of cellular-adhesion programs, consistent with conservation of tumor-immune communication networks across species (**Supplementary Figures 3A-B**). Even though the analyses of end-point spontaneous murine tumor samples captured less overall immune cells compared to available human datasets, their interactions with the murine cancer cells was still predictive of immunosuppression.

**Figure 3:**
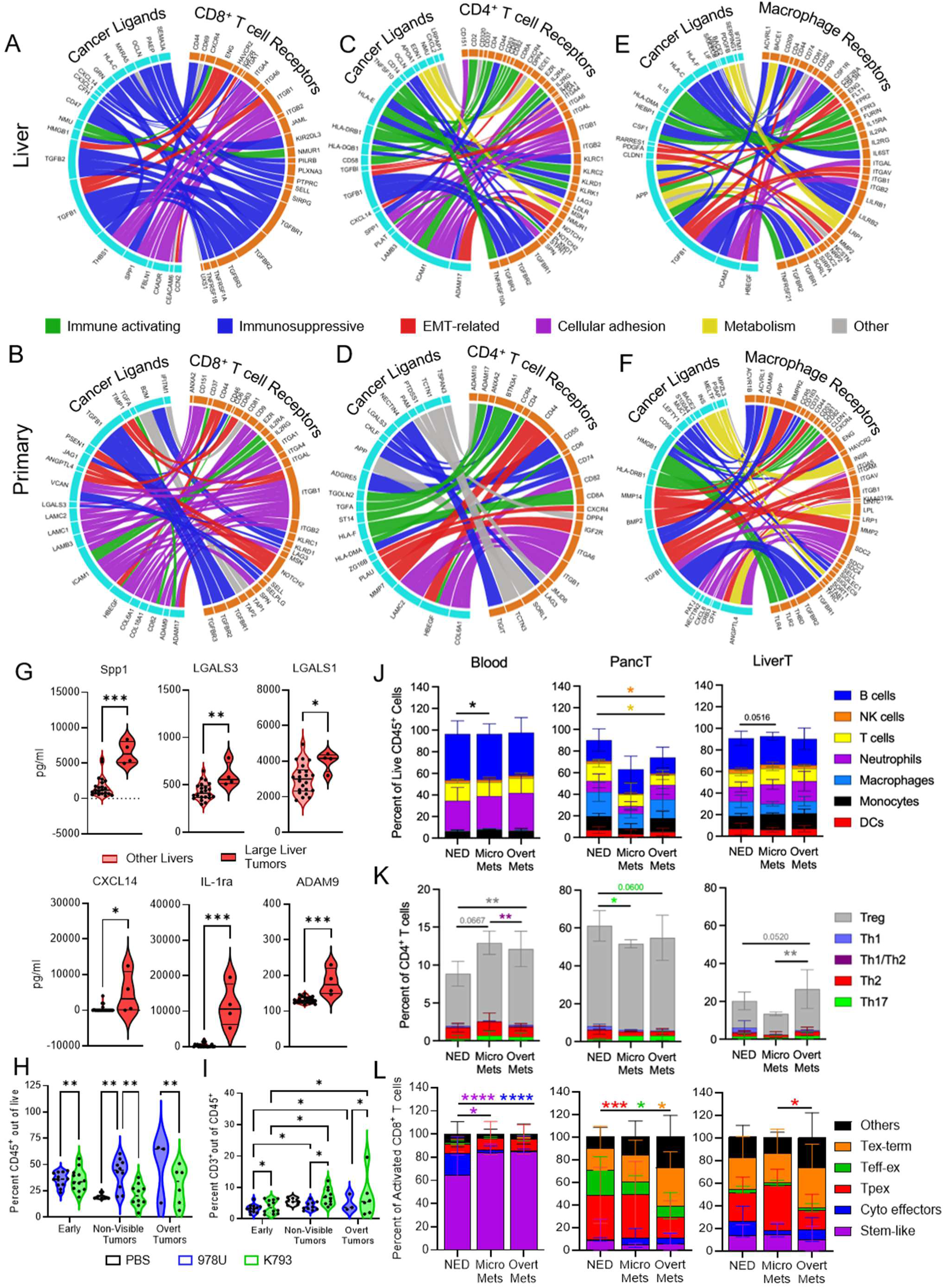
Cross-Species Analyses Reveal Immunomodulatory Signaling in PDAC Liver Metastases. **A-F**. Predicted ligand–receptor interactions between cancer cells and immune populations across human samples. Interactions between cancer cells and CD8⁺ T cells in liver metastases (A) and primary pancreatic tumors (B), cancer cells and CD4⁺ T cells in liver metastases (C) and primary pancreatic tumors (D), and cancer cells and macrophages in liver metastases (E) and primary pancreatic tumors (F). **G**. Violin plots showing the mean expression of ligand-receptor associated proteins identified in liver metastatic cancer cells through NicheNet analyses across liver samples. Mann-Whitney T test. **H-I.** Flow cytometric quantification of CD45⁺ immune populations and CD3^+^ T cells across hemisplenic transplanted livers. Two-way ANOVA. **J.** Flow cytometric quantification of immune cell populations gated sequentially such that “Lin-” denotes exclusion of all lineage markers listed in the preceding populations: B cells (CD19^+^), NK cells (CD19^-^, NK1.1^+^), T cells (CD19^-^, NK1.1^-^, CD11b^-^, CD3^+^), Neutrophils (CD19^-^, NK1.1^-^, CD3^-^, Ly6G^+^), Macrophages (Lin^-^, F4/80^+^), Monocytes (Lin^-^, Ly6C^+^), and Dendritic cells (Lin-, MHC-II^+^, CD11c^+^) Two-way ANOVA. NED, n = 5-12; Micro Mets, n = 3; Overt Mets, n = 13-19. **K.** Flow cytometric quantification of CD4^+^ Lineage^-^ (NK1.1, B220, CD19, Gr1, Ter119, and CD11b) CD3^+^ T cell subsets: Treg (Foxp3^+^), Th1 (Foxp3^-^ Tbet^+^ GATA3^-^), Th1/Th2 (Foxp3^-^ Tbet^+^ GATA3^+^), Th17 (Foxp3^-^, Tbet^-^, GATA3^-^ RORy^+^). Two-way ANOVA. NED, n = 4-10; Micro Mets, n = 2-3; Overt Mets, n = 12-15. **L.** Flow cytometric quantification of activated CD8^+^ T cell (CD69^+^ CD62L^-^) subsets: stem-like (Tcf1^+^), cytotoxic effector (Gzmb^+^ Prf1^+^) progenitor-exhausted (Tpex) (Tcf1^+^ PD-1^+^), intermediary effector-exhausted (Teff-ex) (PD-1^+^ CX3CR1^+^), and terminally exhausted (Tex-term) (PD-1^+^ TIGIT^+^). Two-way ANOVA. NED, n = 10-12; Micro Mets, n = 2-3; Overt Mets, n = 15-19. * p<0.05, ** p<0.01, *** p<0.001, ns, not significant.

Analysis of cancer cell–CD4^+^ T-cell interactions revealed a broader signaling landscape than observed for CD8^+^ T cells. In human datasets, both primary pancreatic tumors and liver metastases demonstrated enrichment of numerous ligand-receptor interactions involving inflammatory, adhesion-associated, and immune-regulatory pathways, consistent with the diverse function of CD4^+^ T cells in PDAC progression and metastasis (**Figures 3C-3D**). In contrast, murine liver metastases were comparatively enriched for immune-activating signaling interactions whereas primary pancreatic tumors demonstrated enrichment of EMT-associated and immunosuppressive signaling programs (**Supplementary Figures 3C-3D**). These findings are consistent with prior studies linking CD4^+^ T-cell signaling to EMT induction, extracellular matrix remodeling, and metastatic progression in PDAC (43–47).

Given the abundance of macrophages within PDAC liver metastases (48–51), we evaluated cancer cell-macrophage communication networks. Both human samples and murine models demonstrated enrichment of immunosuppressive and EMT-associated ligand-receptor interactions in primary pancreatic tumors and liver metastases (**Figures 3E-F** and **Supplementary Figure 3E-F**). Several conserved signaling axes, including TGFB1-, SPP1-, APP-, and integrin-associated interactions, were observed across species, supporting the potential role of macrophage-mediated niche remodeling during metastatic outgrowth.

To determine whether the predicted ligand-receptor interactions were reflected at the protein level, we performed a quantitative mouse cytokine multiplex array on high- (K793) and low- (978U) liver-tropic cell lines, their corresponding liver tumors generated through hemisplenic transplantation, and PBS-injected controls. Samples were collected at two time-points: five days post-surgery to capture early dissemination and humane endpoint to assess established metastatic disease. As expected, the high liver-tropic lines generated overt liver metastases which were microdissected and compared to all other liver samples with no evidence of tumor growth. Hierarchical clustering identified five protein-expression clusters, with clusters 1–3 enriched for liver-derived samples, cluster 4 enriched for cultured tumor cells, and cluster 5 selectively enriched in microdissected overt liver metastases (**Supplementary Figure 3G**). Overt liver metastases had increased protein expression of SPP1, LGALS3, LGALS1, CXCL14, IL-1RA, and ADAM9 relative to other liver-derived samples (**Figure 3G** and **Supplementary Figure 3H**), consistent with the transcriptomic analyses and prior reports of metastatic progression, immune suppression, and extracellular matrix remodeling (52–58). Since the array was performed on bulk tissue, these measurements establish that the predicted ligands are present in the metastatic niche.

To further assess immune infiltration patterns during metastatic progression, we performed flow cytometric analysis of hemisplenic-transplanted livers. Early after dissemination, high- and low-metastatic tumors differed in the proportion of CD45^+^ immune cells; by endpoint, low-metastatic tumors exhibited greater overall immune infiltration than high-metastatic tumors (**Figure 3H**). Within the immune compartment, high-liver-tropic tumors consistently contained a greater proportion of T cells than low-metastatic tumors across time points (**Figure 3I**). Macrophages represented the dominant immune population independent of tumor type and were significantly enriched in tumor-bearing samples relative to time-matched PBS controls, whereas B-cell populations remained comparatively stable across groups (**Supplementary Figures 3I-J**). To determine how metastatic burden influences systemic immune composition, we performed deeper immunological profiling of blood, primary pancreas, and matched liver tumors across multiple dual-transplant models (59–63). All mice had primary tumors and were further stratified according to liver disease status: no evidence of disease (NED), micrometastatic disease, or overt metastases. We observed increased frequencies of circulating and liver tumor-associated monocytes in mice with micrometastatic lesions compared with NED animals. In the primary pancreatic tumors, increasing liver tumor burden also associated with lower frequencies of T cells and NK cells (**Figure 3J**). These data support previous conclusions that metastatic progression is accompanied by systemic immune remodeling (7,64–66).

We next characterized T cell differentiation states. Within the CD4^+^ compartment, Th17 cell frequencies increased in pancreatic tumors with escalating liver metastatic burden. Given the context-dependent roles of Th17 cells in tumor immunity, further studies will be required to determine their contribution to the primary TME in late-stage disease. In contrast, Tregs represented the dominant CD4^+^ subset among those profiled across all tissues and increased in both circulation and liver tumors with increasing metastatic burden. Notably, the composition of circulating CD4^+^ T cells closely tracked that of the liver tumors, supporting the possibility that progressive liver metastatic disease is associated with systemic enrichment of immunosuppression (**Figure 3K**).

Within the activated CD8^+^ T cells, cytotoxic effector cells were significantly reduced in both micrometastatic and overt metastatic disease compared with NED, accompanied by a corresponding increase in the stem-like population. In contrast, the primary pancreatic tumors exhibited progressive remodeling toward dysfunctional states, with significant shifts in progenitor-exhausted (Tpex), intermediary effector-exhausted (Teff-ex), and terminally exhausted (Tex-term) CD8^+^ T cells from NED to overt metastases. Liver metastases displayed a more restricted pattern, with a significant decrease in the Tpex population between micrometastatic and overt metastatic lesions. Notably, intermediary effector-exhausted CD8^+^ T cells were less abundant in liver metastases than in primary pancreatic tumors, consistent with a distinct immune-remodeling of the metastatic microenvironment (**Figure 3L**). Together, these findings suggest that liver metastasis is accompanied by progressive remodeling of the intratumoral CD8^+^ T-cell compartment, while micrometastatic lesions may retain a comparatively less dysfunctional immune state that could provide a window for therapeutic intervention.

Collectively, these analyses indicate a conserved liver-tropic cell state associated with a distinct tumor–immune communication. Cross-species ligand–receptor analyses revealed shared immunoregulatory and matrix-remodeling interactions, including TGFB1, SPP1, and LGALS3, between malignant cells and immune populations, particularly CD8^+^ T cells and macrophages. These interactions were further supported by cytokine profiling, which detected the corresponding proteins within overt metastatic lesions. Cellular profiling in murine models showed that these environments become progressively more immunosuppressive with metastatic outgrowth, with Treg enrichment in liver tumors and circulation and a shift of intratumoral CD8⁺ T cells toward terminal exhaustion. Notably, this remodeling was less advanced in micro-metastatic than in overt lesions, suggesting that the metastatic immune environment is not fixed at seeding but evolves during outgrowth.

### Site-specific immune composition distinguishes pancreatic and liver tumors independently of metastatic phenotype

To determine whether the tumor–immune signaling programs identified above were reflected in site-specific tissue organization, we performed 43-plex immunofluorescence profiling of matched pancreatic tumors, liver tumors, and adjacent normal tissues from dual-transplants of high-tropic (K793 and 7160c2) and low-tropic (O739) cell lines (**Supplementary Table 7**).

Consistent with prior studies, substantial heterogeneity was observed across all tumor microenvironments (67–71) (**Figures 4A**-**B** and **Supplementary Figures 4A-B**). Despite intertumoral variability, liver tumors consistently exhibited greater immune-cell infiltration than pancreatic tumors across all cell lines, with CD45^+^ cells preferentially localized along invasive tumor borders (defined as approximately 35 µm inside the tumor/uninvolved border) (**Figure 4C**) (7,72).

**Figure 4:**
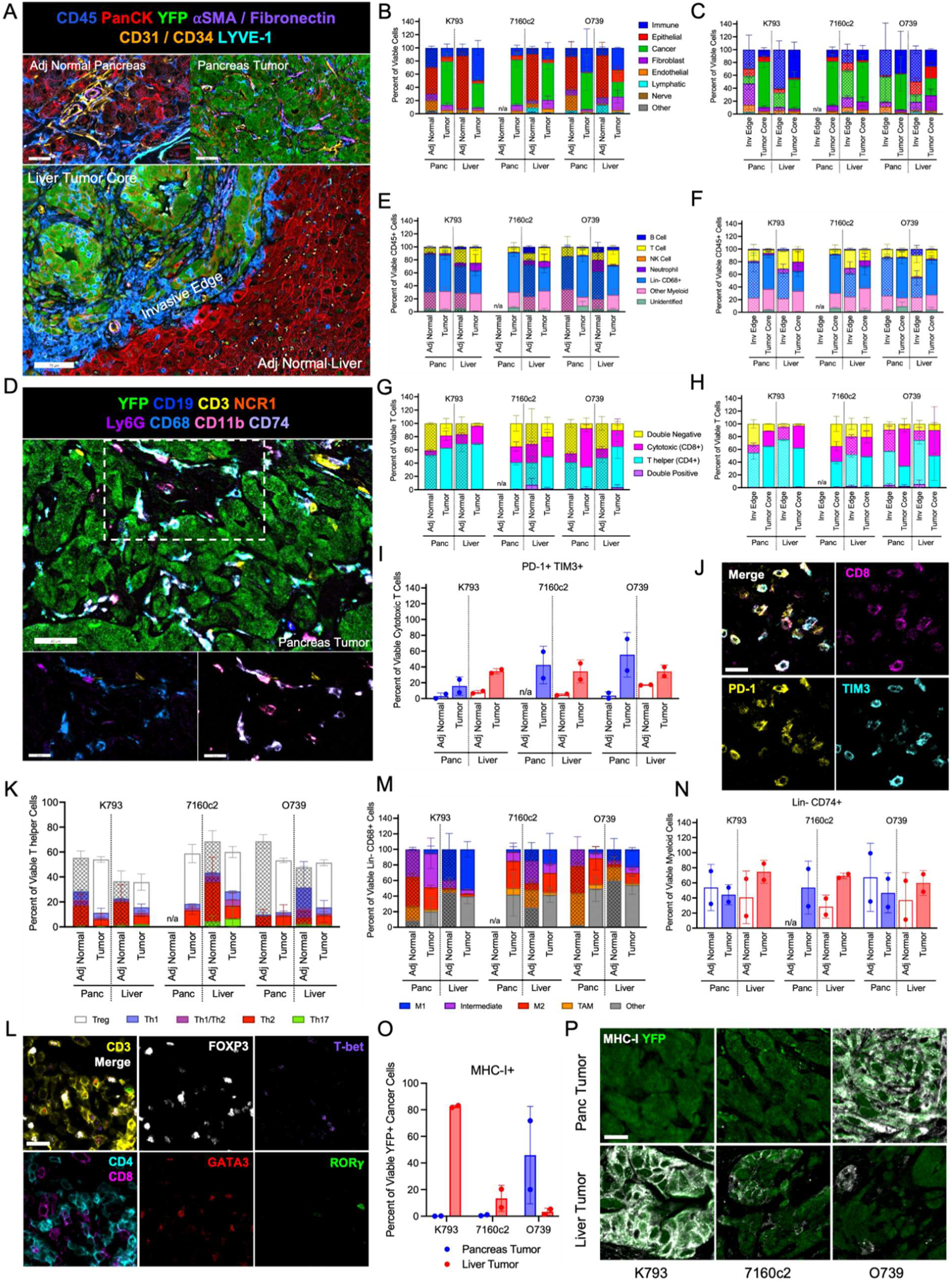
Anatomical Site Is a Major Determinant of PDAC Immune Composition. **A**. Representative multiplex immunofluorescence micrographs (scale 40 µm) of immune, epithelial, cancer, and stromal markers within adjacent normal pancreas and pancreas tumor from K793 cell line, with a representative image (scale 75 µm) of the same markers within K793 liver showing tumor core, invasive edge and adjacent normal tissue. **B-C**. Frequency of Immune (CD45^+^), Epithelial (Pan cytokeratin and/or E-Cadherin^+^), Cancer (YFP^+^), Fibroblast (αSMA^+^, FAP^+^, Desmin^+^, and/or Vimentin^+^), Endothelial (CD31^+^ and/or CD34^+^), Lymphatics (LYVE-1^+^), Nerve (Neurofilament Heavy Chain^+^), or Other (DAPI^+^ but Negative for all other markers) out of viable (cleaved-caspase-3 negative) cells in specified ROIs. **D**. Representative micrographs (scale 40 µm) showing immune cell markers in 7160c2 pancreas tumor. Inset image (scale 20 µm) showing Ly6G, CD68, and CD3 (bottom left) and CD11b, CD74, and CD3 (bottom right). **E-F**. Frequency of immune cell populations B cells (Ly6G^-^ CD19^+^), T cells (Ly6G^-^ CD19^-^ NCR1^-^ CD3^+^), NK cells (Ly6G^-^, CD19^-^, NCR1^+^), Neutrophils (Ly6G^+^), Lin^-^ (Ly6G^-^ CD19^-^ NCR1^-^ CD3^-^) CD68^+^ cells, other myeloid (negative for the previous markers, positive for CD74, CD11b, CD163, CD206, or CD86 in any combination), and unidentified (negative for all markers but CD45^+^) out of viable immune cell populations (as defined in B-C) in specified ROIs. **G-H**. Frequency of T cell subtypes out of viable T cells in specific ROIs. **I**. Quantification of PD-1^+^ TIM3^+^ Cytotoxic T cells. **J**. Representative micrographs of PD1^+^ TIM3^+^ CD8^+^ cells, taken from O739 pancreas tumor (scale 20 µm). **K**. Frequency of T helper cell transcriptional regulators out of viable CD4^+^ T helper cells: Treg (Foxp3^+^), Th1 (Foxp3^-^ Tbet^+^ GATA3^-^) Th1/Th2 (Foxp3^-^ Tbet^+^ GATA3^+^), Th17 (Foxp3^-^, Tbet^-^, GATA3^-^ RORy^+^). **L**. Representative micrographs of T cell subtypes with transcription factors to mark T helper subsets (scale 20 µm). The merge photo does not contain CD4 and CD8 for population clarity. Image taken along O739 pancreas tumor border along a lymph node. **M**. Frequency of macrophage populations within viable Lin^-^ CD68^+^ cells showing M1 markers (CD86^+^ CD206^-^), Intermediate (CD86^+^ CD206^+^), M2 markers (CD86^-^ CD206^+^ CD163^-^), Tumor Associated Macrophages – TAM (CD86^-^ CD206^+^ CD163^+^) and Other (remaining Lin^-^ CD68^+^ immune cells). **N**. Frequency of Lin-CD74^+^ cells within viable myeloid cells. **O.** Frequency of MHC-I expression on viable YFP^+^ cancer cells according to tumor ROIs. **P**. Representative micrographs of MHC-I and YFP expression according to tumor sites with MHC-I quantification (scale 20 µm). All quantifications were done on standardly processed images; brightness and contrast were adjusted for figure visualization only. For all panels, ROIs pooled from n = 2 mice, with liver met ROIs ranging from 1-10 distinct ROIs per mouse.

Myeloid cells, particularly lineage-negative CD68^+^ populations, represented the dominant immune compartment across tumors (**Figures 4D** and **4E**), consistent with flow cytometric analyses and prior studies (73,74). T cells were more frequent in liver tumors than pancreatic tissue, localizing more on the tumor’s invading edge (**Figures 4E** and **4F**), as previously reported (75,76), while the highest percentage of T cells out of all viable cells was observed in the liver tumor of the high-liver tropic line K793 (**Supplementary Figure 4C**). Consistent with previous studies (73), CD4^+^ T cells were the major T cell population in all tumors from the high tropic cell lines, but their liver tumors had more CD8^+^ T cells than their matching primary tumors. CD8^+^ T cells constituted the majority of the low-tropic O739 pancreas tumor T cell population, as its CD4^+^ T cells were more localized at the invasive edge (**Figure 4G** and **4H**). Although some recent patient studies have reported reduced T-cell and myeloid-cell infiltration in liver metastases relative to primary tumors, those analyses primarily compared untreated primary tumors with post-treatment metastatic biopsies (9). The therapy-naïve setting examined here may therefore capture distinct site-specific immune states as supported by other therapy-naïve work (74).

Consistent with the immunosuppressive nature of PDAC, CD8^+^ T-cell exhaustion was elevated in tumors relative to adjacent normal tissues across all models (**Figures 4I** and **4J**). Site-specific differences in exhaustion were variable across cell lines and did not consistently track with PD-L1 expression (**Figures 4I** and **4J** and **Supplementary Figures 4D and 4E**) (42,74). Pro-tumor CD206^+^ macrophages (TAMs), Tregs, and Th2 cells were the predominant immunosuppressive populations across tumors, consistent with previous reports (**Figures 4K-M**) (71,77,78). In contrast, normal liver tissue associated with the low-metastatic O739 line exhibited increased neutrophil, Th1-cell, and exhausted CD8^+^ T-cell frequencies relative to other normal liver samples, suggesting immune features that may influence metastatic permissiveness (**Figure 4E, 4I**, and **4K**).

Furthermore, because immune cell function is closely linked to antigen presentation during TME evolution (79), we quantified cancer cell MHC-I expression and myeloid cell expression of CD74, a chaperone protein for MHC-II antigen presentation. Liver tumors consistently exhibited slightly higher frequencies of Lin^-^ CD74^+^ cells than pancreas tumors across all cell lines (**Figure 4N**), suggesting differences in antigen presentation between anatomical sites that may contribute to the increased immune cell and CD4^+^ T cell infiltration in liver tumors. In the pancreatic tumor, MHC-I expression was highest in the low-liver tropic O739 cells; however, liver tumor MHC-I expression was highest in the high-tropic K793 and 7160c2 cancer cells (**Figures 4O** and **4P**). Together, these findings indicate that antigen-presentation features differ across tumor sites and metastatic phenotypes and may contribute to site-specific tumor–immune interactions independently of CD8^+^ T-cell exhaustion.

Multiple patient and murine studies have highlighted how immune cell localization, high myeloid and Treg frequencies, and T cell dysfunction contribute to immunosuppression in primary and metastatic TMEs (22). Consistent with these observations, our analyses identified increased immune-cell infiltration along tumor invasive edges, elevated frequencies of Lin^-^ CD68^+^ CD206^+^ cells, Tregs, and exhausted CD8^+^ T cells in tumor tissues. Liver TMEs were distinct from pancreatic with greater immune-cell infiltration, altered T cell composition, and enrichment of immunosuppressive cell populations, consistent with observations from human PDAC metastases (7,42,72,74). These findings demonstrate that anatomical site exerts a strong influence on PDAC immune composition, generating distinct immune ecosystems in pancreatic and liver tumors regardless of metastatic phenotype. These observations prompted us to next examine whether tumor–immune spatial organization further distinguishes primary and metastatic disease.

### Spatial analyses identify liver-specific interactions between cancer cells and immunosuppressive T-cell populations

Many cellular functions are regulated across space, making cellular frequencies only one part of functional assessment. We previously showed that the spatial proximity of TME components is associated with patient outcome (80). Therefore, we quantified proximity, infiltration, and clustering between cancer cells and the TME populations across pancreatic and liver tumors to define site-specific tumor–immune architecture and map immunosuppressive niches (81).

Mean minimum distance and G-function analyses demonstrated that multiple CD4^+^ T cell subsets were both positioned closer to and more highly enriched around cancer cells in the liver tumors than the pancreatic tumors (**Figures 5A-5C and Supplementary Figure 5A**), indicating substantial liver-specific differences in cancer cell – T cell spatial organization (82). We also observed significantly more Tregs near exhausted (PD-1^+^ TIM3^+^) CD8^+^ cytotoxic T cells in liver tumors overall (**Figure 5D**), supporting prior reports linking Tregs with CD8^+^ T cell exhaustion in PDAC (83). Interestingly, pancreas tumors from low liver-tropic cell lines showed more CD8^+^ T cells near Lin^−^ CD68^+^ cells compared to pancreas tumors from high liver-tropic cell lines (**Supplementary Figure 5B**), suggesting a potential macrophage–cytotoxic T cell interaction distinct to the low-liver tropic primary tumors (67). Together, these findings identify liver-specific immunosuppressive niches characterized by increased spatial association of Tregs with exhausted CD8^+^ T cells that may promote liver metastatic burden.

**Figure 5:**
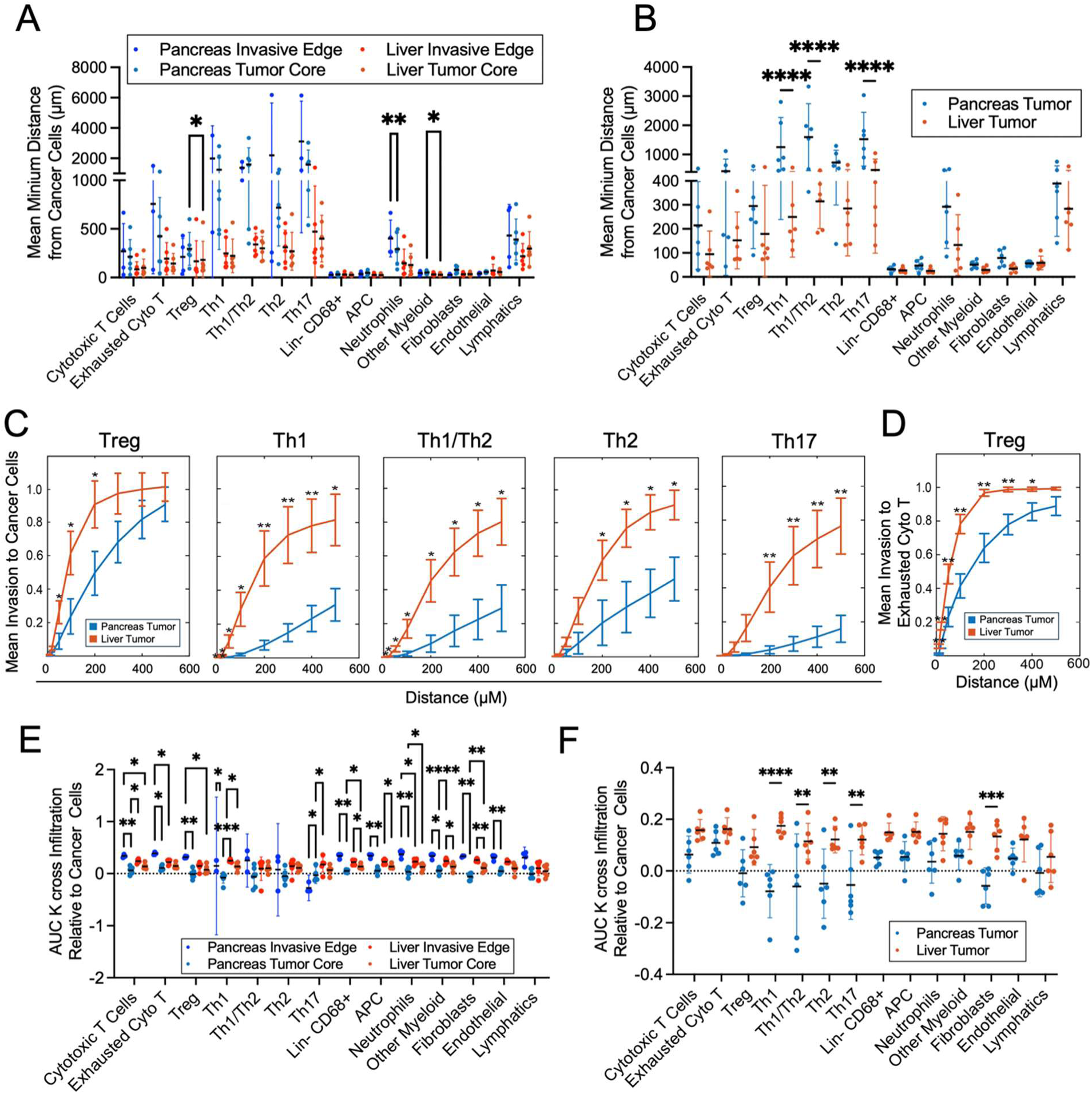
Liver tumors display enhanced TME infiltration around cancer cells. **A-B**. Mean median distance in µm of the indicated TME cell type around cancer cells in the indicated regions of interest. Data are a pooled weighted sum of each ROI by mouse, weighted by the number of cancer cells in each ROI. n = Pancreatic Invasive Edge: 4, All other groups: 6. Mean with 95% Confidence interval. ANOVA with mixed-effects model, matching across mice. **C**. G function plots showing mean invasion to cancer cells within liver and pancreas tumor ROIs by all CD4^+^ T helper cell subsets, where the x-axis is distance from a cancer cell, and the y-axis is probability of the indicated infiltrating cell being within that radius. Two-sample, two-sided t-tests **D**. G function plots showing mean invasion to exhausted (PD-1^+^ TIM3^+^) cytotoxic T cells within liver and pancreas tumor ROIs by Tregs. Two-sample, two-sided t-tests **E-F**. Area under the curve (AUC) of the K-cross function for the pattern of infiltration of the indicated TME cell type around cancer cells in the indicated regions of interest. Data are a pooled weighted sum of each ROI by mouse, weighted by the number of cancer cells in each ROI. n= pancreatic invasive edge: 4, all other groups: 6. Mean with 95% Confidence interval, ANOVA with mixed-effects model, and matching across mice. * p<0.05, ** p<0.01, *** p<0.001, **** p<0.0001, ns, not significant.

To evaluate patterns of cellular clustering, we next performed K-cross function analyses and found a reduction in the clustering between cancer cells and TME components in pancreatic tumor cores, with significant differences across several immune and non-immune populations (**Figure 5E**). Th17 cells at the pancreatic invasive edge were segregated from cancer cells, whereas tumor cores showed intermixed clustering. Comparisons between pancreatic and liver tumors revealed significantly greater clustering between cancer cells and all Foxp3^−^ CD4^+^ T cell subtypes in liver tumors (**Figure 5F**). These data highlight the complex and site-specific ecosystems around cancer cells.

Consistent with prior human studies (9), liver tumors exhibited increased infiltration of T cells, B cells, αSMA^+^ fibroblasts, and macrophages despite no significant differences in their proximity to cancer cells between tumor sites. Our analyses further support previous reports describing liver metastases as localized immunosuppressive niches with T cells enriched at tumor margins and increased Treg infiltration surrounding cancer cells relative to pancreatic tumors (7). Together, ligand–receptor, cytokine, and spatial analyses reveal that liver metastases possess a distinct tumor–immune architecture characterized by enhanced interactions between cancer cells and immunosuppressive T-cell populations. These findings identify spatial organization as a defining feature of PDAC liver metastases and complement the conserved immune signaling programs observed across species.

### T cells constrain PDAC liver metastatic outgrowth

Given the association between T-cell composition and liver metastatic progression observed in both patient studies and our analyses, we next sought to determine whether T cells functionally influence metastatic outgrowth. Low T-cell infiltration within primary PDAC tumors has been associated with increased liver metastatic risk and reduced patient survival (6,80). Consistent with these observations, pancreatic tumors generated in orthotopic and dual transplants of the high liver-tropic cell lines contained fewer T cells than pancreatic tumors in low liver-tropic line transplants (**Figure 6A and Supplementary Figure 6A**). To independently evaluate this relationship, we utilized two previously characterized KPC-derived cell lines reported to show low (6419c5) or high (6620c1) T cell infiltration within orthotopically transplanted tumors (84). While both cell lines efficiently formed pancreatic tumors, only the low T cell line 6419c5 developed large liver tumors that were within the range of the established high liver-tropic tumors (**Figures 6B** and **6C**). These findings further support an inverse relationship between T cell infiltration in the primary tumor and liver metastatic outgrowth.

**Figure 6:**
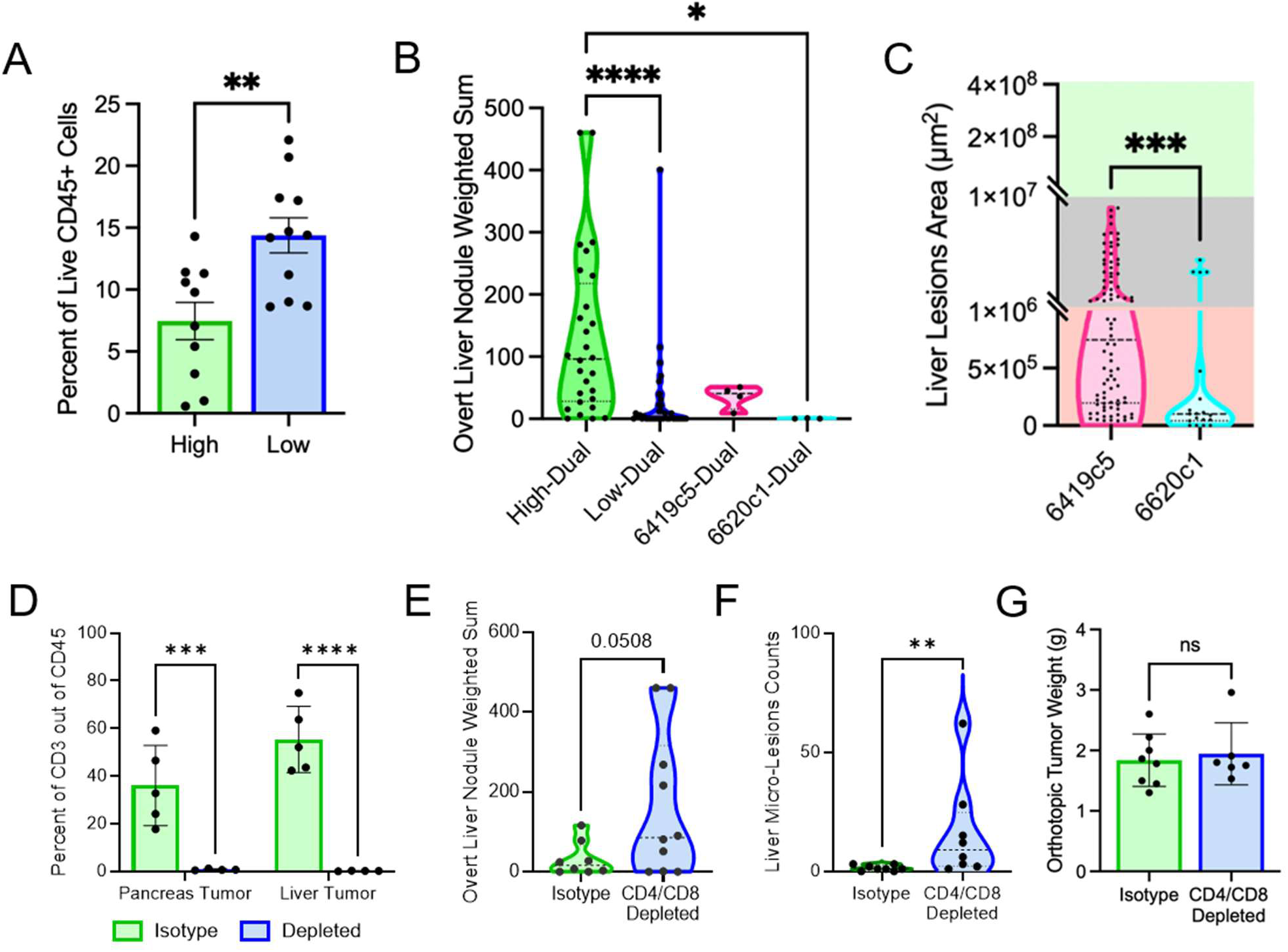
T cells constrain PDAC liver metastatic outgrowth. **A.** Flow cytometric quantification of CD3^+^ T cell in the pancreas tumors from dual transplants. n = High lines: 10, Low lines: 11. Welch’s t test. **B**. Weighted sum by size of overt liver nodules of dual injected. n = High lines: 28, Low lines: 38, 6419c5: 4, 6620c1: 3. Kruskal-Wallis One-Way ANOVA. **C.** Quantification of each metastatic liver nodule area of the indicated cell lines. Axis breaks correspond to clinical imaging groups of micrometastases (red), borderline detectable (gray), and detectable (green). Kruskal-Wallis One-Way ANOVA. **D**. Flow cytometric quantification of CD3^+^ T cell in the pancreas or liver tissue of dual transplanted mice treated with isotype or anti-CD4 and anti-CD8 depleting antibodies. n= Isotype: 5, anti-CD4/CD8: 4. Matching Two-Way ANOVA with Sidak’s Multiple comparison tests. **E**. Weighted sum by size of overt liver nodules of dual-injected 978U and 2838c3 lines treated with isotype or anti-CD4 and anti-CD8 depleting antibodies. n= Isotype: 8, anti-CD4/CD8: 10. Welch’s t-Test. **F**. Count of micrometastatic liver nodules. n = Isotype: 8, anti-CD4/CD8: 8. Mann Whitney test. **G.** Weights of orthotopic tumors from dual transplant. n= Isotype: 8, anti-CD4/CD8: 6. Mann Whitney test. * p<0.05, ** p<0.01, *** p<0.001, **** p<0.0001, ns, not significant.

The enrichment of immunosuppressive CD4^+^ T cell populations, exhausted CD8^+^ T cells, and suppressive tumor–T cell spatial niches in liver metastases prompted us to investigate whether T cells directly contribute to constraining metastatic burden. To test this, mice bearing low liver-tropic, high T-cell tumors were treated with CD4- and CD8-depleting antibodies in the dual-transplant system (**Supplementary Figure 6B**). Combined T-cell depletion resulted in significantly increased overt and microscopic liver metastatic burden compared with isotype-treated controls, while primary pancreatic tumor size was unaffected (**Figures 6D-G**).

Together, these findings provide functional support for the association between T-cell abundance and liver metastatic progression observed in both clinical datasets and our experimental models. Although these studies do not identify the specific T-cell subsets or mechanisms responsible for restricting metastatic growth, they demonstrate that T-cell-mediated immune surveillance contributes to limiting PDAC liver metastatic outgrowth. These results further support the biological relevance of the tumor–immune interactions and spatial immune features associated with PDAC liver metastases.

## Discussion

Liver metastasis remains the primary driver of PDAC mortality at diagnosis and recurrence. Although several studies have advanced our understanding of PDAC metastatic progression (10–13), the biological features that distinguish liver metastases from primary pancreatic tumors remain incompletely defined, and experimental systems capable of evaluating matched primary and metastatic disease are limited. Here, we developed a syngeneic dual-transplant model system that generates matched pancreatic and liver tumors within the same host and leveraged this platform to identify biological features associated with PDAC liver metastatic outgrowth. Across murine and human datasets, we identified a conserved metastatic cell state enriched in liver metastases and associated with poor patient outcome. In parallel, we found that liver metastases exhibit distinct tumor–immune communication networks, altered immune composition, and unique spatial organization characterized by enhanced interactions between cancer cells and immunosuppressive T-cell populations. Finally, functional T-cell depletion studies provided evidence that T-cell-mediated immune surveillance contributes to constraining liver metastatic outgrowth. Together, these findings support the concept that conserved metastatic cell states and spatial immune remodeling are defining features of PDAC liver metastases.

Using several primary tumor-derived KPC cell lines, we show that while all lines efficiently form pancreatic tumors, their capacity for liver outgrowth varies widely despite shared oncogenic driver mutations. These findings further support the idea that cancer cell-intrinsic programs contribute to metastatic progression and interact with the liver microenvironment to govern metastatic colonization and outgrowth, which prior studies have left underappreciated (85). Comparative transcriptomic analyses identified a high liver-tropic transcriptional program that consistently marked metastatic cancer cells across multiple murine and human datasets. Importantly, this signature was enriched in PDAC liver metastatic lesions, detectable in circulating tumor cells from patients with metastatic disease, associated with reduced overall survival, and absent from liver metastases arising from breast and colorectal cancers as well as hepatocellular carcinoma. The enrichment of this program within early disseminated tumor cell populations further suggests that it captures biological processes associated with metastatic seeding and early colonization (27–30). Although prospective validation is required, these observations raise the possibility that this metastatic cell state could serve as a biomarker to detect liver-disseminated disease below the threshold of clinical imaging.

While the high liver-tropic metastatic state identified here was consistently associated with liver metastatic progression across species and datasets, the functional contributions of specific genes and pathways within this program remain to be mechanistically defined. Prior studies have linked metabolic fitness, mitochondrial adaptation, and cellular stress responses to successful metastatic colonization (27–31); our analyses support these concepts by identifying enrichment of metabolic programs within highly liver-tropic tumors alongside reduced inflammatory and stress-associated signaling. Future mechanistic studies will be necessary to determine whether these transcriptional features directly contribute to dissemination, colonization, or metastatic persistence within the liver.

Our integrated analyses further revealed extensive remodeling of tumor–immune communication within liver metastases. Cross-species ligand–receptor analyses identified enrichment of predicted TGFB1-, SPP1-, and LGALS3-associated signaling networks between cancer cells and immune populations, while multiplex protein profiling demonstrated increased abundance of several corresponding immunomodulatory proteins within overt liver metastases. Together, these findings suggest the existence of conserved tumor–immune communication programs associated with metastatic progression. However, these observations remain correlative, and pathway-specific perturbation studies will be required to determine whether particular signaling axes directly contribute to immune suppression, metastatic niche formation, or tumor outgrowth within the liver.

Beyond signaling interactions, multiplex spatial profiling demonstrated that anatomical site strongly shapes the immune composition of PDAC tumors. Consistent with previous studies of metastatic disease (7,67,73,74,78), liver metastases exhibited distinct immune ecosystems characterized by increased immune-cell infiltration, altered T cell composition, and enrichment of immunosuppressive populations, including Tregs, Th2 cells, and CD206^+^ Lin^-^ CD68^+^ cells. While these populations have each been implicated in PDAC progression and immune dysfunction, our data suggests that the collective organization of these cells within the metastatic microenvironment may be equally important. Of note, our analyses did not include the murine macrophage-selective marker F4/80. Although CD68 is a commonly used macrophage marker, some overlap with monocytes and dendritic cells is possible, and future studies incorporating additional lineage markers will be important for refining the precise identities of these myeloid populations. Still, our data support claims that myeloid cells and T cells play important roles in maintaining and promoting tumor development (86–90).

A particularly notable finding was the distinct spatial organization of TME cells within liver metastases. Multiple CD4^+^ T cell subsets, fibroblasts, and antigen-presenting cells were positioned closer to cancer cells and displayed greater clustering around tumor cells in liver metastases than in primary pancreatic tumors. In addition, Tregs were more frequently associated with exhausted CD8^+^ T cells. These results support previous studies linking Tregs, myeloid cells, and fibroblasts to CD8 cell exhaustion and dysfunction in the TME (83,91,92). These observations suggest that liver metastases contain localized immunosuppressive niches that differ substantially from those present in primary pancreatic tumors. Because spatial relationships can influence cellular communication, antigen presentation, and immune function independent of cellular abundance alone, these findings highlight spatial immune remodeling as an important component of metastatic progression. Future studies will be needed to determine whether these spatial interactions directly regulate T cell dysfunction or represent downstream consequences of metastatic adaptation within the liver.

To begin addressing the functional relevance of these immune observations, we evaluated the relationship between T cells and metastatic outgrowth. Consistent with clinical studies showing an association between reduced T cell infiltration and liver metastatic progression (6,93), cell lines characterized by lower T cell abundance generated greater liver metastatic burden. Furthermore, depletion of CD4^+^ and CD8^+^ T cells preferentially increased liver metastatic outgrowth while having minimal effects on primary tumor growth. Although these experiments do not define which T cell subsets are responsible for restricting metastatic progression or how their function becomes impaired within liver metastases, they provide functional support for the hypothesis that T cell-mediated immune surveillance contributes to limiting PDAC liver metastatic outgrowth. These findings also complement our spatial analyses by demonstrating that immune populations identified within metastatic lesions are not merely associated with, but functionally linked to, metastatic burden.

Several limitations should be considered when interpreting these findings. While the dual-transplant system enables the generation of matched pancreatic and liver tumors and preserves cell line-specific metastatic phenotypes, it bypasses early steps of the metastatic cascade, including local invasion and intravasation. Similarly, the ligand–receptor analyses presented here are predictive in nature, and the T cell depletion studies broadly target lymphocyte populations rather than specific immune subsets. Consequently, the mechanisms linking metastatic cell states, immune signaling, and spatial immune architecture remain incompletely resolved. Nonetheless, the ability to generate matched primary and metastatic tumors within an immunocompetent setting provides a tractable platform for addressing these questions in future studies.

## Conclusion

Collectively, our findings identify a conserved metastatic cell state and a distinct spatially organized immune ecosystem that characterize PDAC liver metastases across murine models and human disease. By integrating matched primary and metastatic tumors with transcriptomic, proteomic, and spatial analyses, we provide evidence that liver metastases are distinguished not only by cancer cell-intrinsic programs but also by unique patterns of tumor–immune communication and organization. These results establish a framework for investigating dynamic, site-specific mechanisms of metastatic progression and nominate tumor cell states, immune signaling networks, and spatial immune interactions as candidate vulnerabilities in PDAC liver metastasis.

## Methods

### Cell lines

Six primary pancreatic tumor cell lines were derived from the KPCY mouse model (Pdx1-Cre; LSL-Kras^G12D^; P53^R172H/+^; R26-LSL-EYFP), which closely recapitulates the genetic landscape and progression of human pancreatic ductal adenocarcinoma (PDAC). Two lines serve as EMT controls. The 28B line was derived from an adenovirus expressing Cre into the pancreas of a KP-Cdh1^F/F^ mouse, resulting in pancreatic cancer cells with E-cadherin (Cdh1) knockout (mesenchymal-stabilized), and K793 was derived from a KPCY mouse with dual Snail1 and Twist1 knockout (Snai1^F/F^; Twist1^F/F^) (epithelial-stabilized)(12). Two additional KPCY-derived tumor cell lines, 978U and O739, were established from near-cousin parental KPCY mice to the EMT-modified lines. All four of these lines were isolated from colonies with a mixed genetic background that was approximately 50-60% C57Bl/6J, genotyping for MHC B-haplotype showed all lines to be homozygous for the B haplotype corresponding to C57Bl/6 or 129J backcrossed line. The remaining lines, 2838c3, 7160c2, 6419c5 and 6620c1, from the laboratory of Ben Z. Stanger, MD, PhD, University of Pennsylvania, were purchased from Kerafast and were reported to be fully backcrossed at least 10 generations onto the C57Bl/6 line (Supplementary **Table 1**) (84). All cell lines were cultured in RPMI-1640 medium supplemented with 20% fetal bovine serum (FBS Genesee Scientific 25-550H) and 1% antibiotic/Antimycotic (Fisher Scientific MT30004CI) under standard conditions (37°C, 5% CO_2_).

Doubling times were determined by plating 100,000 cells in 6 well plates in triplicate and counting every 12 hours for 108 hours. The doubling time was determined by calculating the slope of the line y = mx+c.

### Transplants

All mice were housed under standard housing conditions at the University of Alabama at Birmingham (UAB) animal facilities. Investigators were not blinded for group allocation but were blinded for the assessment of the phenotypic outcome in histological analyses. All open body cavity surgeries were performed using aseptic technique. Mice were anesthetized (isoflurane) to the point of areflexia to toe pinch. The surgical site was prepared by removing fur around the incision site with a commercially available depilatory cream (Nair™ Hair Removal Lotion for Sensitive Skin; Church & Dwight Co., Inc., Ewing, NJ, USA), containing calcium thioglycolate as the active depilatory agent. After removing any gross contamination with a simple wipe of the skin with clean water, chlorhexidine is applied in three alternating sets of application with sterile alcohol. A left paramedian skin incision with surgical scissors was given on the abdomen, and a plane was separated between the subcutaneous tissue and peritoneum, then an incision through the peritoneum. Next, the pancreas and spleen were surgically exteriorized. For orthotopic injections: syngeneic mouse pancreatic tumor cell lines (1x10^4^-1x10^5^) in 25 µL of sterile PBS using a Hamilton syringe (Fisher Scientific NC2156196, 50 µL volume with blunt end for soft tissue injections) were directly injected into the pancreatic tail. Hemisplenic injections were performed as previously described (18,23). Brief modifications are as follows: the spleen was exposed and delivered through the peritoneum and clamped in between the pancreatic veins using a hemostat, and then bifurcated using a high-temperature cautery pen (Fisher Scientific NC9074124) on the dorsal side of the hemostat. The dorsal hemispleen was allowed to return to the body cavity, while maintaining hold of the hemostat and ventral hemispleen, which was then injected with 100µl of cell suspension (1x10^4^, 1x10^5^, or 5x10^5^) followed by 150µl of PBS pulled previously in the same 0.5 cc insulin syringe, keeping the syringe upright at all times to avoid mixing the cell and the “chaser” phases. After injection and partial resolution of blanching, the ventral hemispleen was removed by cauterizing the draining pancreatic veins. Portal vein injections were performed as previously described (17). We injected 1x10^5^ in 25 µL using ½ cc insulin syringe, followed by gentle pressure with a hemostatic gauze (Fisher Scientific 50 223 2753) to achieve hemostasis.

For T cell depletion studies, mice received intraperitoneal administration of either isotype control or depleting antibodies beginning 72 hours prior to tumor transplantation and again 72 hours following surgery. Antibodies were subsequently re-administered every 72 hours until experimental endpoint, defined by the development of large tumor burden. Mice received 200 µg each of InVivoMAb anti-mouse CD4 (Bio X Cell, Cat# BE0003-1-R050mg) and anti-mouse CD8 (Bio X Cell, Cat# BE0061-R050mg) antibodies, or 200 µg of rat IgG2b isotype control, anti-keyhole limpet hemocyanin (Bio X Cell, Cat# BE0090-R050mg). All antibodies were prepared in the manufacturer-recommended InVivoPure dilution buffer. T cell depletion was validated at endpoint (**Figure 6E**).

### GEMM

Characterization of disease progression and genotyping for the *Pdx1-Cre; LSL-Kras^G12D^; P53^R172H/+^ ; R26-LSL-EYFP* (herein referred to as KPC or KC if wildtype for *P53^R172H/+^*) mice were previously described (94). The mice were maintained on a mixed background, bred according to Mendelian ratios, and developed normally (data not shown). Males and females were utilized equally. No randomization method was used, and all mice of the desired genotype were enrolled in the study. Age, disease progression, sex, and quantified metrics are reported for each mouse in the Supplementary **Table 8**.

### Overt Metastasis Scores

Upon necropsy, the presence or absence of visible metastatic nodules was assessed in the liver, lung, diaphragm, peritoneum, mesentery, gut tissues, spleen, and surgical incision. The liver was additionally quantified based on an agreed-upon binning system for the relative size of the nodules, later measured to be approximately the indicated areas: tiny (0.785 mm^2^), small (3.561 mm^2^), medium (6.242 mm^2^), large (25.95 mm^2^), full lobe (NA). The count of the nodules in each category was multiplied by an increasing factor of three over the previous bin and summed for the weighted overt liver lesion score except for full lobes, where a scaling factor was selected that allowed 4 full liver lobes to be the highest value while avoiding excess score inflation (tiny*1 + small *3 + medium *9 + large * 27+ full *115).

### Histology

Histology and histopathological scoring were performed similarly to those previously described (12). In brief, tissues were collected and fixed for 24-48 hours at room temperature in 10% neutral buffered formalin (Fisher Scientific 22-050-104). Formalin-fixed tissues were embedded in paraffin and sectioned at 5 μm thickness. Sections were stained with hematoxylin and eosin (H&E; Fisher Scientific 22-050-306, 22-050-116, and 22-050-110). Histopathological measurements were made by scoring H&E-stained tumors for relative percentages of each histopathological phenotype: normal (non-neoplastic), PanIN/ADM, Adenocarcinoma, which was a combined score of well-differentiated, moderately-differentiated, and poorly-differentiated PDAC, Sarcomatoid carcinoma, or necrosis. Data presented as the percentage of viable (non-necrotic) tumor (non-normal) tissue. When tumor histology was missing or of poor quality, the mice were excluded from all histological analyses, and this was determined blinded from group allocation. Microscopic tumors in the liver were observed in H&E-stained tissue sections at four depths, 150 μm apart. A summary incidence of the presence of any metastasis for each mouse is presented in Supplementary **Table 8**.

Metastatic lesion areas were processed using Visiopharm Oncotopics Analysis software, assisted with a custom-generated AI user-trained app on each tissue section using scanned 4x images. All calls were manually validated as correct. The scans were generated on the Echo Revolution microscope and exported as tif images. These exported tif images were then converted to a pyramidal ome.tiff format with LZW compression using the bftools package in Windows PowerShell using the following prompt: “bfconvert -tilex 512 -tiley 512 -noflat -pyramid-resolutions 3 -pyramid-scale 2 -compression LZW”. The user-generated Visiopharm apps were generated on the following parameters: 1. Tissue Finder_DeepLearning_AI app identifying background (glass), Tissue using default UNet settings at 1x magnification, and trained for 1000+ iterations across several representative slide scans. Post-processing was performed to fill holes and smooth outlines. Tissue ROIs were manually reclassified as pancreas, liver, lung, or diaphragm as present in the slide. Slide scans with large areas of adjacent primary tumor were manually modified to clear those areas from the tissue ROI. All calls were manually checked and corrected if needed. MetsFinder_Deeplearning_AI app identified Tumor and Normal classifications restricted to the “Target ROI” either Liver or Lung tissues as appropriate. Trained at 4x magnification using default UNet deeplearning for approximately 500,000 (liver) and 120,000 (lung) iterations with post-processing to clean up label areas to normal or tumor. All labels were manually checked for accuracy of calls and adjusted accordingly. Data outputs included nodule counts, total nodule area, total target tissue area, percent tumor burden (total nodule area/ tissue area), and individual nodule area and maximum diameter. As tissues were used in many different assays and sample preparations, the area of microscopic lesions are plotted individually.

Immunofluorescence staining was performed on 5 µm thick sections that were deparaffinized and rehydrated. Heat-mediated antigen retrieval was performed in Tris-EDTA with 0.1% Tween 20 (pH 9.0) for 15 minutes at 95°C in the EZ Retriever microwave, BioGenex. After cooling in dH2O, the tissues were outlined with pap pen (Fisher Scientific NC1710579). All wash steps were performed with TBS and blocking/primary antibody diluent were performed in 1% BSA (Fisher AAJ65097A1) with 3% goat serum (Fisher Scientific 1CN19135680) in TBS. Antibody staining was performed in a primary cocktail of Gata6 (1:500, Cell Signaling 5851S) and GFP (1:500, Aves Labs 5851S) followed by a secondary cocktail of Goat anti-Chicken-AlexaFluor 488 (1:1000, Invitrogen A11039) and Goat-anti-Rabbit-CoraLite594 (1:500, ProteinTech A11039) followed by EpCam-Cy5 (1:200, Abcam ab237385), and finally counter stained with DAPI (1:20,000 Fisher Scientific D1306 resuspended in dimethylformamide) and mounted with VectaShield mounting media. Five representative images per tumor were taken with the Echo Revolution using the 20x objective, exported as a tif and processed as previously described into ome.tiff format. These images were quantified using the Visiopharm analysis software for cell segmentation and marker positivity with the Phenoplex guided workflow.

### Immunohistochemistry

FFPE sections of each cell line were deparaffinized and underwent antigen retrieval in Tris-EDTA buffer (pH 9.0) at 95°C for 15 minutes using the Biogenex EZ Retriever system. Slides were then incubated with 3% H_2_O_2_ for 15 minutes at room temperature then washed with TBS. At room temperature, sections were blocked with 1% BSA (Fisher AAJ65097A1) in TBS for 30 minutes, incubated with anti-CD3 (1:200, Abcam, ab11089) for 1 hour, washed with TBS, then incubated with Polink 1 HRP Rat (Orgene, Cat: D35-6) for 30 minutes. ImmPACT DAB eqV (Vector SK-4103-100) was added to each section for 7 minutes then washed with water. Hematoxylin (Fisher Scientific 22-050-306) was used as counterstain. Multiple 10x images were captured for each slide with Echo Revolution microscope, exported as tif images, and analyzed with Visiopharm Oncotopics Analysis software. Nuclei Detection, AI (Brightfield), CD3_IHC_quant185 app identified nuclei and quantified DAB positivity to output the percent CD3^+^ cells per image.

### Bulk RNAseq

Bulk RNA sequencing was performed on previously published KPC-derived PDAC cell lines 978U, K793, O739, 28B, 7160c2, and 2838c3 (12,95). Total RNA was isolated using the TRIzol reagent according to the manufacturer’s recommendations (Fisher Scientific 15-596-026) and submitted to the UAB Genomics Core Laboratory for library preparation and sequencing. Libraries were sequenced on an Illumina NovaSeq 6000 platform using paired-end 100 bp reads, with an average sequencing depth of approximately 69 million reads per sample. FASTQ files were generated using Illumina bcl-convert. Raw sequencing reads were imported into Partek Flow (v12.7.0) and aligned to the mm38 mouse reference genome using the STAR aligner (v2.7.8a). Following post-alignment quality control, gene-level read counts were generated using featureCounts. Data were normalized, and differential expression analysis was performed using ANOVA, with significantly altered genes defined as those with P < 0.05. EMT signature was quantified using z-score normalization across samples, and heatmap values ranged from −2.16 to 4.01. To identify a high liver-tropic gene signature, bulk RNA-seq data from two highly liver-tropic and three low liver-tropic KPC-derived PDAC cell lines were analyzed. Genes were first pre-filtered for *p-value* < 0.05, after which z-scores were calculated across all five cell lines. Genes commonly upregulated in both highly liver-tropic lines, with z-score values > 1.2 exclusively in both highly liver-tropic group, were included in the signature, resulting in a 462-gene high liver-tropic signature. Using biomaRt, mouse genes in the signature were mapped to their human orthologs by querying Ensembl and converting mouse gene identifiers to the corresponding human orthologous gene symbols (96). Pathway enrichment analysis was conducted using Metascape to identify enriched signaling pathways associated with tumor-intrinsic transcriptional programs in these PDAC cell lines.

### scRNAseq analysis

#### Mouse

Publicly available single-cell RNA sequencing (scRNA-seq) datasets were obtained from GSE165534 (12), which profiled whole pancreatic tumor tissue from six KPC mice (three early and three late stage) and six KPC;ST mice (one early and five late stage), as well as liver tissue from two early-stage KPC, three late-stage KPC and four late-stage KPC;ST mice. In the original study, tissues were dissociated into single-cell suspensions and processed for scRNA-seq library generation and sequencing as described. For the present study, scRNAseq FASTQ files were aligned to the mouse reference genome version (mm39) in 10x Genomics CellRanger (version8.0.1).

The acquired gene expression matrices were then loaded into RStudio (version 2023.12.1-402) and analyzed using Seurat version 5 workflow (97). Quality control filtering retained cells with 200–7,000 detected genes, 500–50,000 UMIs, and <10% mitochondrial RNA content. The *SCTransform()* function was applied to normalize, scale, and identify highly variable features, regressing out mitochondrial fraction. Principal component analysis was conducted with the *RunPCA()* function, and 40 principal components were selected for downstream analyses based on the Elbow plot. Cell clustering was performed with the *FindNeighbor()* and *FindCluster()* functions and a resolution of 1.2, and the clusters were then projected into the Unifold-Manifold Approximation and Projection space with the *RunUMAP()* function. Coarse cell types were manually annotated using canonical gene markers curated from the literature. Finer-grained annotations of T/NK and Myeloid cells were subsequently performed by sub-setting each coarse cell population into smaller Seurat object, repeating the bioinformatics workflow of *SCTransform*, dimensionality reduction, and clustering analyses. In all cases, manual cell type annotations were confirmed using the *FindAllMarkers()* function. Generated Seurat objects have been deposited in a Zenodo notebook available upon publication.

#### Human

The human datasets comprised raw scRNA-seq FASTQ files from 12 independent studies that profiled tumor and immune cells using the 10x Genomics platform. Across studies, samples included primary tumors and associated metastasized tumor microenvironments generated from human PDAC patients (33). For the present study, the processed Seurat object was obtained from Zenodo (https://zenodo.org/records/14199536) and imported into RStudio for downstream re-analysis. More specifically, the original dataset provided coarse cell-type labels, so we performed fine-grained re-annotation of T/NK and Myeloid cell populations. Similar to the mouse dataset, we first subset each compartment into an independent Seurat object and repeated the standard processing pipeline and parameters aligned with the data’s original publication: normalization, identification of highly variable features, scaling, batch correction with Harmony package version 1.2.3 (98), dimensionality reduction, and clustering. Subclusters were first annotated utilizing canonical markers curated from the literature and were verified using Seurat’s *FindAllMarkers()* function.

#### Pseudobulk

Single-cell RNA sequencing data from Loveless et al., 2025 (33) was used to generate pseudobulk expression profiles for downstream analysis. Cells were assigned to their respective biological samples using the *Name* field (renamed as *Sample_ID*). Cells were further classified into biological conditions based on clinical annotations: cells labeled as *Primary tumor* were designated as “Primary,” while cells annotated as *Metastatic lesion* with a metastatic site of *Liver* were designated as “Liver.” Pseudobulk profiles were generated by summing raw gene expression counts across all cells belonging to each Sample_ID and condition using the AggregateExpression() function in Seurat. This resulted in a gene-by-sample count matrix, where each column represents the aggregated transcriptomic profile of a given sample. The pseudobulk count matrix was subsequently treated as bulk RNA-seq data and normalized using the trimmed mean of M-values (TMM) method implemented in edgeR to account for differences in library size and compositional bias across samples. Normalized counts were then transformed to log2 counts per million (logCPM) using the cpm() function with a prior count of 1 to stabilize variance and mitigate the influence of lowly expressed genes. The resulting logCPM matrix was used for downstream gene set variation analysis.

#### Gene Set Variation Analysis (GSVA)

GSVA package was used to quantify pathway activity at the sample level (99). Gene expression matrices, including both pseudo-bulked single-cell RNA-seq data and bulk RNA-seq datasets, were first normalized and log-transformed prior to analysis. A curated liver metastasis gene signature was used as input, and gene identifiers were matched to the corresponding expression matrices to ensure compatibility. Enrichment scores were computed independently for each sample based on the ranked expression of genes, enabling robust estimation of pathway activity without reliance on cross-sample distribution assumptions. This approach was selected to accommodate heterogeneity across datasets, including differences in study origin, sample composition, and sequencing platforms. The resulting scores were used for downstream comparative analyses across biological conditions and cohorts. For non-pseudobulked single-cell and spatial transcriptomics datasets, we applied an orthogonal Seurat-built-in algorithm, *AddModuleScore()*, to quantify gene set expression levels per cell or per spot.

#### NicheNet

We comparatively analyzed across species the intra- and inter-cellular communication patterns between cancer and immune cell populations in both the primary and liver metastatic tumor microenvironments by applying the NicheNet (v2.1.0) algorithm on the aforementioned human and mouse scRNA-seq datasets (100). We followed the workflow recommended by the developers in their recent Nature Protocols publication (101), implementing both the sender-agnostic and sender-focused approaches.

For each analyzed “receiver” cell population, we first performed differential expression analysis across microenvironmental sites and retained site-upregulated genes representing the population’s site-specific transcriptional program. Differentially expressed genes (DEGs) were required to be expressed in at least 5% of cells within the population, meet a minimum log fold-change threshold, and pass multiple-testing correction (adjusted p ≤ 0.05). We used the default log fold-change cutoff of 0.25 as per the Protocol in most analyses, with the only exception of a more stringent cutoff of 1 for the sender-agnostic analysis in the human dataset to mitigate detection of small-effect changes arising from substantially larger cell numbers compared to the mouse dataset. After differential expression analysis, we subsetted the original human and mouse Seurat objects into two separate objects representing the primary tumor and liver metastatic microenvironments, retaining only cells in each corresponding site before performing the sender agnostic approach to ensure that the inferred ligand-receptor interactions are site-specific. All genes detected in the dataset were used as background genes.

Implementing the sender-focused approach, we then constructed a site-specific list of ligands by identifying ligand genes expressed in sender cell populations with at least one of their cognate receptors expressed in receiver cell populations, using cells derived separately from either the primary pancreatic or liver metastatic tumor microenvironment. The selected ligand and receptor genes must be present in at least 5% of the respective cell population. Based on the receiver population’s differentially expressed gene program, we performed NicheNet ligand activity analysis to prioritize ligands whose predicted downstream targets best explain the observed transcriptional changes in the receiver. We then further filter out the top 20 ligands and their associated top 200 target genes, calculate the ligand-target regulatory potential score, and visualize the interactions within the top 25th percentile as similar to default parameters in the Protocol. Subsequently, we pinpoint receptors in receiver cells that interact with the identified ligands and construct ligand-receptor heatmaps. Determination of immune suppressive or immune activating interactions was manually annotated based on literature searches. Circos plots were generated in R using the circlize package to visualize ligand-receptor interactions between cancer cells and immune cell populations. Interaction pairs identified through ligand-receptor analysis were grouped based on their reported biological functions into five major categories: immunosuppressive, immune activating, EMT-related, cellular-adhesion related, metabolism-related, and other interactions. Functional categorization was determined through literature-supported annotation of the known roles of ligands, receptors, and associated signaling pathways. Interaction weights derived from the ligand-receptor analysis were used to scale connection strength within the Circos plots.

### ELISA Array

Cell-line pellets and minced tumor tissue samples were snap frozen in 1.5 mL microcentrifuge tubes in liquid nitrogen and shipped to RayBiotech Inc. for protein isolation and analysis with the Mouse Cytokine Array Q400 (Catalog # QAM-CAA-400-1), which produces quantified protein concentrations for 400 proteins primarily associated with immune signaling. Raw data was processed for the dilution factor used for the cell line pellets, normalized with a z-score, and plotted as an interactive heatmap with hierarchical clustering and sub-groupings with k means value of 5 using R and the ComplexHeatmap library. Differential Expression analysis was run on dilution adjusted values with R using the limma library. Of the ligand genes identified in the transcriptomic ligand–receptor interaction studies, 26 corresponding proteins were represented within the Mouse Cytokine Array Q400 panel and were included for comparative downstream analyses.

### Spatial transcriptomic quantification

#### Spatial Transcriptomics Data Processing

Visium spatial transcriptomics data were processed using Seurat (v4.4.0) in R (v4.4.1). Spot-level gene expression was normalized using SCTransform variance-stabilizing transformation. Cancer-associated spatial ecotypes were identified using ISCHIA clustering (cc_ischia_10 resolution), and tumor-dominant ecotypes (CC2, CC3) were confirmed independently in Liver and Pancreas (primary tumor) tissue subsets using RCTD-based cell-type deconvolution (Robust Cell Type Decomposition), which provided estimated proportions for 15 cell populations per spot.

#### Signature Scoring

Two independently derived gene signatures were scored: a 460-gene High-Liver-Tropic signature and a 47-gene Low-Liver-Tropic signature (Supplementary Tables 5 and 6, respectively, of the associated manuscript). Human gene symbols were matched against the Visium SCT assay feature set; 400 of 425 human-mapped High-Tropic genes (94.1%) and 36 of 47 Low-Tropic genes (76.6%) were detected and used for scoring. Signature scores were computed per spot using Seurat::AddModuleScore() on SCT-normalized data, applied to the full dataset prior to subsetting to the cancer-ecotype cohort.

#### Local Spatial Statistics

Local immune and stromal cell-type enrichment was quantified using the Getis-Ord Gi* statistic (Getis & Ord, 1992, Geographical Analysis, 24(3), 189–206), computed independently for each tissue section using the Spatial neighbor graphs were constructed using k = 6 nearest neighbors (knearneigh/knn2nb), with each spot’s own neighborhood including itself

(include.self()) to compute the Gi* (starred) form, using row-standardized binary spatial weights (nb2listw, style = "B") and localG() with zero.policy = TRUE. Gi* was computed using the full tissue section (all spots, not restricted to the cancer-ecotype subset) to preserve accurate spatial neighborhood context.

#### Statistical Modeling

Associations between signature score and local Gi* enrichment were tested using linear mixed-effects models fit with the — lme4 and lmerTest package. The model specification was: Gi* ∼ signature_score + tumor_epithelial_proportion + (1 | patient), fit by REML. Tumor epithelial cell proportion (from RCTD deconvolution) was included as a covariate to control for a compositional confound (signature score correlated with tumor purity: Liver rho = 0.35–0.42 across cohorts and signatures for High-Tropic; near-zero to weakly negative for Low-Tropic). Patient was modeled as a random intercept to account for unequal per-patient spot representation, following identification of a Simpson’s paradox effect in an uncorrected pooled analysis. P-values were adjusted for multiple comparisons across the 10 tested cell types using the Benjamini-Hochberg method.

#### Independent Validation — Published GSEA-Based Scores

As an independent validation not dependent on RCTD deconvolution, local immune/stromal signal was additionally assessed using 29 published Functional Gene Expression Signatures (FGES; Bagaev et al., 2021, Cancer Cell) scored using pre-computed, spot-level GSEA-based scores from prior work by this group. For each patient, cancer-ecotype spots were divided into tertiles by signature score; the mean FGES score in the top versus bottom tertile was computed per patient and averaged (unweighted) across patients, analogous to the random-intercept correction used in the primary mixed-effects analysis.

#### Site Comparison

The full analysis pipeline (ecotype identification, signature scoring, Gi* computation, mixed-effects modeling, and FGES validation) was independently repeated for cancer-ecotype spots from Pancreas (primary tumor) tissue, using the identical ISCHIA ecotype definition and statistical corrections, to test for site-specific versus conserved associations between signature score and local immune/stromal composition.

#### Spatial Visualization

Spatial figures were generated using Python with matplotlib (v3.10.8), scipy (v1.17.1), and pandas. Smoothed signature-score and cell-type enrichment surfaces were generated by cubic interpolation (scipy.interpolate.griddata) onto a regular grid, masked to exclude extrapolated regions beyond three times the median nearest-neighbor spacing from any observed spot, and rendered as filled contour plots (matplotlib.pyplot.contourf).

### Software and Package Versions Confirmed

- R v4.4.1
- Seurat v4.4.0
- SeuratObject v4.1.4
- dplyr v1.1.4
- tidyr v1.3.1
- readxl v1.4.5
- knitr v1.50
- Matrix v1.7.3
- Python v3.12.3 (figure generation environment)
- matplotlib v3.10.8
- scipy v1.17.1
- pandas v3.0.2
- numpy v2.4.4
- adjustText v1.4.0

### Spatial proteomic analysis

#### Multiplex immunofluorescence staining

The UAB Flow Cytometry and Single Cell core’s Lunaphore COMET seqIF staining platform was used to perform cyclical multiplex immunofluorescent staining on FFPE (Formalin fixed, paraffin embedded) mouse dual transplant tumor slides. Antigen retrieval was performed at 107° C for 15 minutes using the Biogenex EZ Retriever system and EZ-AR2 Elegance (BioGenex HK547-XAK) buffer. Primary antibodies were diluted in multi-staining buffer (BU06, Lunaphore) **Supplementary Table 7,** incubating for 4 minutes while secondary antibodies were diluted in antibody diluent (LI-COR # 927-65001) **Supplementary Table 7** and DAPI counterstain, incubating for 2 minutes. Image buffer (BU09, Lunaphore) was used with DAPI, TRITC, and Cy5 during imaging with exposure times of 25 ms, 250 ms, and 400 ms, respectively. A 2-minute elution step was performed at 37° C after each cycle using elution buffer (BU07-L, Lunaphore) followed by a 30-second quenching step with quenching buffer (BU08-L, Lunaphore). Upon completion of imaging, an aligned, stitched, flat-field corrected 12-bit OME-TIFF image file is generated; then, autofluorescence subtraction in each channel is performed using the Lunaphore Horizon Viewer. Processed images were exported post-LZW compression as pyramidal OME.TIFFs for additional analysis.

#### Spatial proteomic image quantification and positivity

COMET images were analyzed using Visiopharm Oncotopix image analysis software. ROIs were manually drawn around each tissue of interest, including pancreatic tumor, adjacent normal pancreas tissue, pancreas tumor invading edge, adjacent normal liver, liver nodule, and liver nodule invasive edges. Each ROI was denoted by a mouse identifier and enumerated within the tissue. Notable areas of immune infiltration were analyzed in their own ROI as lymph nodes, potential tertiary lymphoid structures, or general myeloid aggregates according to the presence of lymph vessels (LYVE-1^+^ staining) and germinal centers (organization of CD19 and CD3 stains). Tumor ROIs were assigned based on tissue morphology, YFP staining, and vimentin expression. Invasive edge ROIs were standardized by eroding tumor ROIs 125 pixels (about 35 µm) then manually adjusting them to include surrounding CD45^+^ cell clusters slightly beyond that limit. Cell segmentation was performed with an AI nuclei detection app based on the DAPI channel, using CD31, CD45, and Pan cytokeratin, and α-SMA as cell membrane defining markers. Apps, Visiopharm overlays, and images are provided in the Zenodo notebook upon publication.

Once cells were identified, specific populations were phenotyped according to marker expression and percent positivity within the label object. Thresholds for marker expression were specific for each slide as each cell line and tissue type had different autofluorescence values and epitope expression. The following cell labels were made according to specific marker combinations: Lymphatics, Endothelial, Immune cells, Epithelial, Nerve Fibers, Cancer cell, Neutrophils, B cells, T cells, NK cells, CD74^+^, CD74^+^ CD11c^+^, CD74^-^ Ly6C^+^, Other CD11b^+^, T helper cells, Cytotoxic T cells, T regs, Th17, Th1, Th2, Th1/Th17, Th1/Th2, Treg/Th17, Treg/Th1, Fibroblasts, Hepatocytes (epithelial cells in liver ROIs), and CAF (fibroblasts in tumor and invasive edge ROIs). Other labels were made as post-processing intermediates to allow phenotyping specific to certain tissues on each slide. For example, with O739 tissue, differing densities in the immune cells required object percentages and thresholds to be different between tumors, lymphoid tissue, and normal pancreas and liver tissue for CD45, Ly6G, CD19, CD3, and CD74. For 7160c2, CD45, E-cadherin, and pan cytokeratin thresholds were set according to tumor and normal tissue. For K793, CD45, YFP, E-cadherin, and pan cytokeratin thresholds were set according to tissue, and in the normal mouse tissue microarray, CD45 positivity was set differently for normal pancreas and normal liver, as the normal pancreas had a much higher background. Since CD19 is a more specific immune population marker than CD3, CD19^+^ cells were isolated before CD3 to allow CD3^+^ cells to be pulled from CD19^-^ Ly6G^-^ CD45^+^ immune cells. However, in the lymph node ROI on O739, CD19^+^ B cells were changed to CD3^+^ T cells if CD3 expression was notably brighter than CD3 expression in the other tissue. This is to account for the tight localization of B and T cells seen in lymph nodes, multiple cells within the 5 µm plane, and imperfect cell segmentation, including pixels from bright CD19 cells next to CD3 cells. Likewise, due to close CD4 CD8 T cell positioning in the lymph node ROI, a similar process was performed on CD4^+^ T helper cells to add more CD8^+^ cytotoxic T cells. Certain immune cells also had specific thresholds in areas of certain ROIs due to high background staining of specific markers. For all slides, cancer cells and hepatocytes were changed to endothelial if CD31 expression was present to account for any CD31 that may have been covered by YFP and epithelial markers when calling those labels first. CD31^+^ cells were not identified before these markers because of the high background in normal pancreas tissue. Because all cells in the normal pancreas ROIs reflected normal morphology, any cancer cell labels identified in normal pancreas ROIs were changed back to the normal cells label because the YFP antibody had high background in normal pancreas tissues.

Object outputs contained form factor, multiplexing identities, ROI, and X and Y values for all cell labels. For multiplexing identities, applied object percentages were added for relevant markers, so downstream data would match the phenotyping App label outputs. Phenoplex Guided Workflow was then used to set the positivity of all markers. Markers used in phenotyping apps were given the same thresholds in the Phenoplex Guided Workflow as were set in the preceding apps. If a marker had more than one threshold, the lower or average of the two settings was applied to best reflect the stain and reduce false positives and negatives. All remaining thresholds were set based on stain expression. Final outputs, including those from the phenotyping app and positivities from the Phenoplex Guided Workflow, were analyzed in Microsoft Excel (Version 2603 Build 16.0.19822.20086) and additional cell phenotypes (Lin^-^ CD68^+^, M1/M2/TAM subtypes, Other Immune, PD-1^+^ TIM-3^+^ cytotoxic T cells, and CAF populations) were assigned based off of binary outputs from the Phenoplex Guided Workflow.

#### Quantification and Pooling of Spatial Interaction Metrics

Using the SPIAT package (81), mean minimum distance summarizes cell–cell proximity by calculating the distance between cancer cells and its nearest neighboring cell population followed by averaging these minimum distances across all cells. The cross K function quantifies cell colocalization by measuring whether a cell population are found more or less frequently around cancer cells with increasing radii. The area under the cross K function curve (AUC) provides a single summary statistic of overall attraction (positive AUC) or repulsion (negative AUC) between cancer cells and target cell population. For each ROI, the mean minimum distance between cancer cells and each annotated cell type, and the cross K function area under the curve (K cross AUC) values quantifying spatial association between cancer cells and each target cell population.

To account for differences in tumor content across ROIs, linear weighting was applied based on the number of cancer cells present in each ROI. For every ROI, the number of viable cancer cells was obtained from the *“Total Cell Counts of Each Label after cleaved caspase-3 removal”* table in the corresponding analysis output and used as a linear weight for subsequent pooling with assistance from an institutional license of M365 Copilot for efficiency of data processing. Pooling was performed with individual mice using two different approaches: (1) pooling by mouse but retaining sub-tissue ROI to produce a value for each ROI: Pancreas invasive edge, Pancreas tumor core, Liver invasive edge; Liver tumor core; and (2) pooling by mouse but only retaining tissue location to produce a value ROI for: Pancreas tumor and Liver tumor. Pooling ROI-level metrics was performed using a weighted mean:

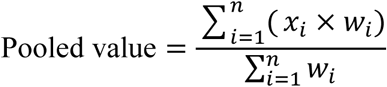

where:

*x*_i_is the ROI-level metric (mean minimum distance or Kcross AUC),
*w*_i_is the corresponding cancer cell count for that ROI, and
*n*is the number of ROIs contributing non-missing values for that mouse and comparison.

ROIs with missing (NA) metric values were excluded from both the numerator and denominator. For transparency, the number of contributing ROIs (*n*) was tracked for every pooled value.

#### G-function

Neighborhood cell-cell colocalization was evaluated with G-functions that measure the percentage of host cells colocalized with a target cell type at a given radius (102,103).The G-function was computed with in-house software developed under MATLAB 2025b and plotted at radius of 10, 20, 50, 100, 200, 300, 400, 500 µm. Specific cell types were chosen to be evaluated with G-function plots under a specifically selected reference/target cell types of comparisons. Significance was determined with two-sample, two-sided t-tests at each radii.

### Statistical Analysis

Statistical analyses were performed using unpaired or paired two-tailed t tests, one-way and two-way ANOVA with multiple comparisons test using GraphPad Prism version 10, as stipulated in the figure legends and detailed source data supplied in the Zenodo notebook upon publication. Chi-square analyses, using SPSS version 31.0 statistical software, were performed comparing indicated groups across multiple gross pathological and histopathological parameters for phenotype incidence. Results are outlined in **Supplementary Tables 3 and 4**, generating values can be found in **Supplementary Table 8**. Kaplan-Meier plots were drawn for time till large tumor, moribundcy, or death of transplanted tumors, and the log-rank Mantel-Cox test was used to evaluate statistical differences, using GraphPad Prism 10. Data met the assumptions of each statistical test, where variance was not equal (determined by an F-test) Welch’s correction for unequal variances was applied.

### Survival Analysis of TCGA-PAAD

Overall survival analysis was performed using the TCGA-PAAD cohort by stratifying patients based on quartiles of the high liver-tropic gene signature score using the Xena-Browser signature tool. Patients in the upper and lower quartiles were compared, and Kaplan–Meier survival curves were generated to evaluate associations between signature enrichment and patient survival outcomes.

### Flow Cytometry

Pancreas and liver tumors/tissues were enzymatically digested to generate single-cell suspensions, which were subsequently filtered through a 70 µm cell strainer and neutralized with complete media (RPMI with 20% FBS and 1% PSA). Red blood cells were lysed using ACK lysis buffer (Gibco™ ACK Lysing Buffer, Catalog #A1049201), followed by washing and cell counting. For flow cytometry staining, 2-3 × 10^6^ cells were incubated with fluorophore-conjugated antibodies against surface markers (**Supplementary Tables 9-13**). Following surface staining and washes, cells for panels with intranuclear markers were fixed and permeabilized using commercially available fixation/permeabilization buffers (eBioscience™ Foxp3 / Transcription Factor Staining Buffer Set, Catalog #00-5523-00) according to the manufacturer’s instructions, followed by intracellular and/or intranuclear antibody staining. After final washes, samples were acquired on the 5-laser Agilent NovoCyte Opteon Spectral Flow Cytometer and analyzed using FlowJo v10.10.1 software according to specified gating strategies (**Supplementary Tables 9-13**)> Dead cells were excluded using viability dye staining, and unstained, fluorescence minus one (FMO), and isotype controls were included where applicable.

### List of Abbreviations

PDAC: Pancreatic ductal adenocarcinoma
TME: Tumor microenvironment
KPC: Kras- and p53-mutant-driven mouse model
GEMM: Genetically engineered mouse models
EMT: Epithelial-to-mesenchymal transition
HS: Hemisplenic transplants
Ortho: Orthotopic (pancreas) transplant
CTC: Circulating tumor cell
FGES: Functional gene expression signatures
NED: No evidence of disease (in the liver)
Tpex: Progenitor-exhausted
Teff-ex: Intermediary effector-exhausted
Tex-term: Terminally exhausted
TAMs: Tumor Associated Macrophages
UAB: The University of Alabama at Birmingham

### Tables List

Supplementary Table 1. Cell line characteristics.

Supplementary Table 2. EMT gene signature

Supplementary Table 3. Overt Incidences at Necropsy by mouse ChiSquare Analysis Results

Supplementary Table 4. Microscopic Incidences by mouse ChiSquare Analysis Results

Supplementary Table 5. High Liver-Tropic Gene Signature

Supplementary Table 6. Low Liver-Tropic Gene Signature

Supplementary Table 7. COMET Antibody Staining Panel

Supplementary Table 8. Mouse Cohorts

Supplementary Table 9. Flow Cytometry Antibody Panel / Gating – T cell, B cell, Myeloid

Supplementary Table 10. Flow Cytometry Antibody Panel / Gating – Myeloid/Lymphoid

Supplementary Table 11. Flow Cytometry Antibody Panel / Gating – CD4 Subsets

Supplementary Table 12. Flow Cytometry Antibody Panel / Gating – CD8 Cell Status

Supplementary Table 13. Flow Cytometry Antibody Panel / Gating – T cell Depletion

## Declarations

### Ethics approval and consent to participate

All animal procedures were reviewed and approved by the UAB Institutional Animal Care and Use Committee (IACUC). All mice will be housed in the animal research facility at UAB. All animals are maintained under the supervision and care of the veterinarians associated with the Animal Facilities at UAB. All facilities are fully accredited by the AAALAC and meet NIH standards as set forth in the “Guide for Care and Use of Laboratory Animals” (DHHS).

### Availability of Data and Materials

Publicly available datasets used in this study included human PDAC single-cell RNA-seq datasets (104), human rapid autopsy RNA-seq datasets (34) and circulating tumor cell RNA-seq data from healthy donors and PDAC patients with liver metastasis(GSE40174) (35) . Bulk transcriptomic datasets of patients with other primary tumors and liver metastasis included breast cancer (GSE110590 and GSE193103) (105,106), colon cancer (GSE81558 and GSE255163) (107,108) and hepatocellular Carcinoma (GSE76427) (109). The other datasets supporting the conclusions of this article are included within the article and its additional files available in Zenodo

upon publication. Bulk RNA-seq FASTQ files generated from the PDAC KPC cell-lines used in this study will be deposited in the GEO database.

### Author Contributions

Ay.Ma. helped design the experimental strategy, performed mouse experiments, conducted flow cytometry on tumor and blood samples, and carried out cell culture studies. She also analyzed transcriptomic and multi-omics datasets, curated publicly available datasets, participated in data interpretation, analysis and visualization, and led writing of the original manuscript draft. C.R.L. helped design the experimental strategy, performed mouse experiments, conducted flow cytometry on tumor and blood samples, carried out histological staining and analyses, collected data, and conducted spatial image analyses. She also participated in data interpretation, data analysis and visualization, and led writing of the original manuscript draft. J.B. carried out histological analyses, including tissue collection, sectioning, staining, imaging, and downstream histological analysis of mouse datasets generated in the lab. T.T. assisted with transcriptomic analyses of mouse and human datasets and assisted in writing the original draft. S.R. analyzed spatial transcriptomic and proteomic data and assisted in writing the original draft. Y.D.F. generated spatial datasets, performed downstream analyses, and contributed to writing the original draft. R.L. assisted with spatial data analysis. M.P. performed cell culture experiments and led bulk RNA-seq data generation and analysis and also reviewed and edited the manuscript. C.R. Assisted in cell segmentation app creation, conceptual support for immunological phenotypes, and reviewed and edited the manuscript. R.S.W. helped design flow cytometry experiments and reviewed and edited the manuscript. A.M.K. analyzed spatial transcriptomic data and assisted with data interpretation.

A.S.M. reviewed and edited the manuscript. S.A. supervised spatial omics analyses, guided data visualization, and reviewed and edited the manuscript. J.L.C. conceived and designed the overall study, provided intellectual direction, supervised the project, guided data analysis and visualization, and contributed to writing and revising the manuscript.

## Acknowledgements

Special thanks to Brody Keys, who assisted in the initial optimizations of the portal vein and orthotopic injections. Special thanks to Harish Pal at the Flow Cytometry and Single Core for his expertise in COMET imaging (supported by the Center for AIDS Research, AI027767, The O’Neal Comprehensive Cancer Center, CA013148, and Shared instrument grant S10OD032296). Special thanks to the Heflin Center of Genomic Sciences Core for their expertise in performing RNA-seq (supported by Shared Instrument Grant 1S10OD032422-01). Special thanks to Brandy Williams at the Comparative Pathology Lab for her expertise in tissue processing and embedding. Special thanks to Justin Major, our Visiopharm Professional Services Representative, for his expert help with novel Visiopharm app design and troubleshooting.

---

**Supplementary Figure 1:**
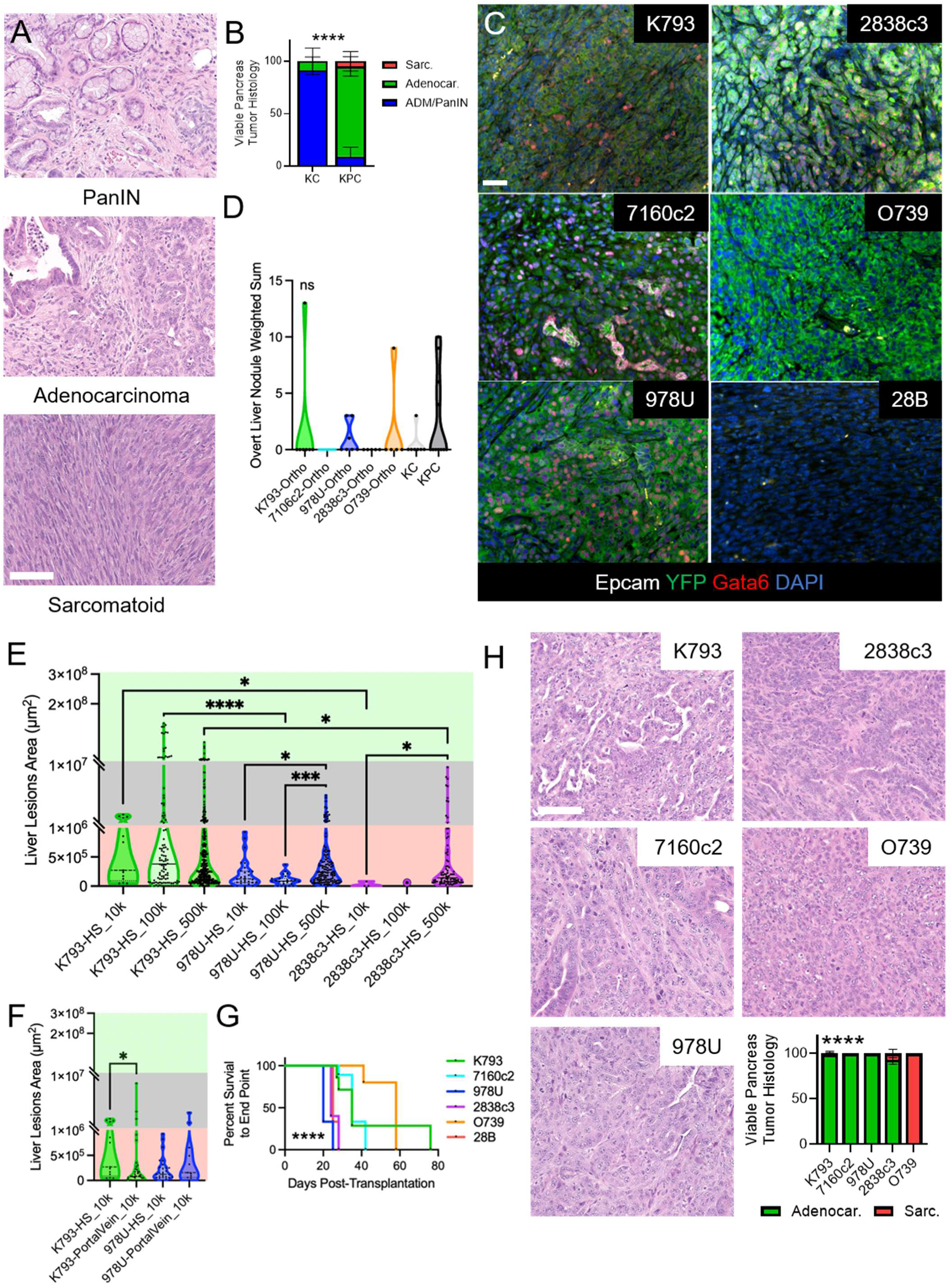
Cancer cell-intrinsic differences define liver metastatic capacity. **A**. Representative histopathology micrographs from spontaneous mouse models. Scale 100 µm. **B**. Relative percentage of histological subtypes in the viable pancreatic cancer regions from the spontaneous KC and KPC mouse models. n = KC: 7, KPC: 10 mice. Two-way ANOVA. Multiple comparisons showed significant differences in PanIN and Adenocarcinoma between the genotypes. Variation of penetrance between the sarcomatoid resulted in non-significance. **C.** Representative micrographs of Epcam, YFP, and Gata6 immunolabeling of orthotopic tumors. Scale 20 µm. Note 28B is a YFP^-^ cell line. **D**. Weighted sum by binned size of overt liver nodules of orthotopically injected (ortho). n = K793:7, 7160c2: 9, 978U: 7, 2838c3: 5, O739: 5, KC: 7, KPC: 10 mice. **E-F**. Quantification of each metastatic liver nodule area of hemisplenic (F) or direct portal vein (G) transplanted cell lines. Axis breaks correspond to clinical imaging groups of micrometastases (red), borderline detectable (gray), and detectable (green). **G**. Percent survival to endpoint of orthotopic transplants. n =K793:7, 7160c2: 9, 978U: 6, 2838c3: 5, O739: 5, 28B: 3 mice. Mantel-Cox test. **H**. Representative micrographs and quantification of histopathology of pancreatic tumors in dual transplants and relative percentage of histological type in the viable pancreatic cancer regions of dual transplanted tumors. n = K793:9, 7160c2: 6, 978U: 6, 2838c3: 5, O739: 6. Two-way ANOVA. Multiple comparisons showed each of the predominantly adenocarcinoma lines was significantly different from O739. Significance determined by Kruskal-Wallis One-Way ANOVA, unless indicated otherwise. * p<0.05, ** p<0.01, *** p<0.001, **** p<0.0001, ns, not significant.

**Supplementary Figure 2:**
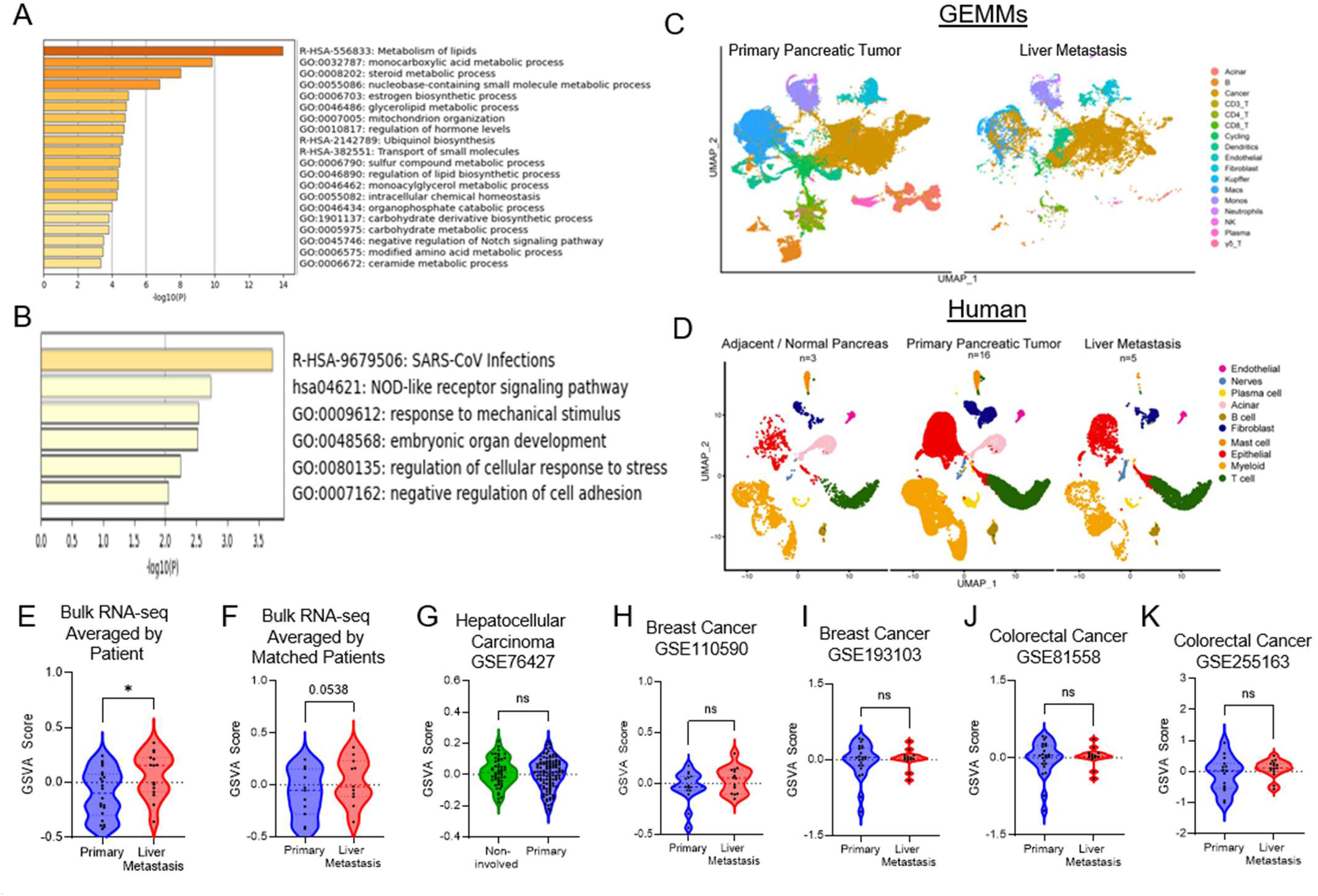
A Conserved Metastatic Cell State Is Enriched in PDAC Liver Metastases. **A-B**. Metascape functional enrichment analysis of the commonly upregulated genes in the high-liver-tropic (A) and low liver-tropic (B) gene signatures. **C**. UMAP visualization of scRNA-seq data showing major cell populations across GEMM. **D.** UMAP visualization of scRNA-seq data showing major cell populations across human PDAC samples. **E-F**. GSVA score of the High-Tropic signature in bulk RNA-seq of all samples from rapid autopsy, averaged by patient (E) and showing only matched primary and liver metastases (F). Unpaired T test or Paired T test, respectively. **G-K**. GSVA score of the High-Tropic signature in bulk RNA-seq of Hepatocellular Carcinoma dataset GSE76427 of matched non-involved liver and the primary (liver) tumor (G), Breast cancer datasets GSE110590 (H) and GSE193103 (I), Colorectal cancer datasets GSE81558 (J) and GSE255163 (K). Unpaired T test with Welch’s correction (G, H and I). * p<0.05, *** p<0.001, **** p<0.0001, ns, not significant.

**Supplementary Figure 3:**
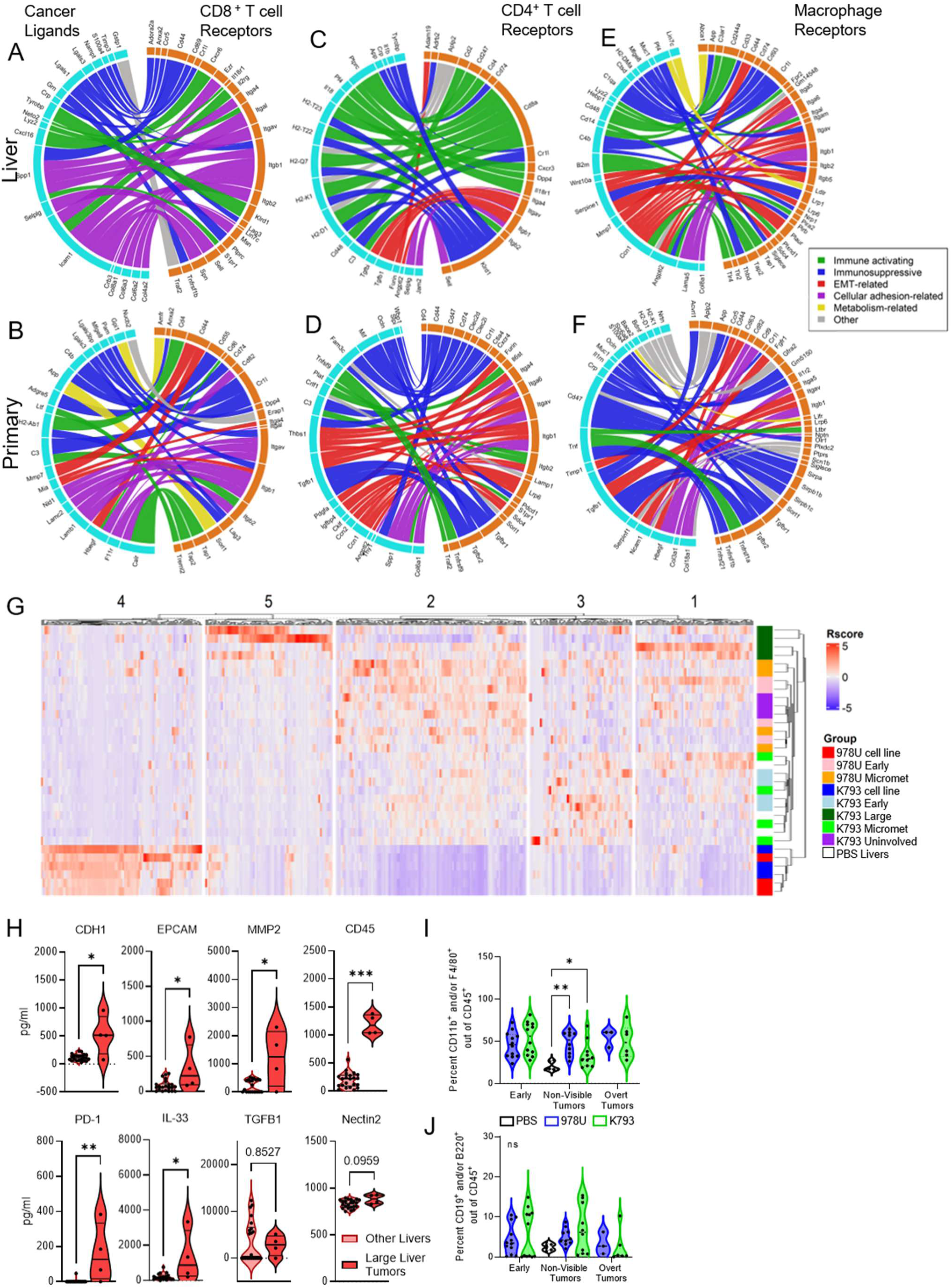
Cross-Species Analyses Reveal Immunomodulatory Signaling in PDAC Liver Metastases. **A-F**. Predicted ligand–receptor interactions between cancer cells and immune populations across PDAC GEMM samples. Interactions between cancer cells and CD8⁺ T cells in liver metastases (A) and primary pancreatic tumors (B), cancer cells and CD4⁺ T cells in liver metastases (C) and primary pancreatic tumors (D), and cancer cells and macrophages in liver metastases (E) and primary pancreatic tumors (F). **G**. Unbiased hierarchical clustering heatmap of cell lines and tumor samples based on a murine cytokine array, with k-means clustering. **H.** Violin plots showing the mean expression of ligand-receptor associated proteins identified in liver metastatic cancer cells through NicheNet analyses across liver samples. Mann-Whitney t test. **I-J**. Flow cytometric quantification of CD11b^+^ and/or F4/80^+^ macrophages (I), and B cells (J) out of viable CD45^+^ cells across hemisplenic transplanted liver. Two-way ANOVA. * p<0.05, ** p<0.01, *** p<0.001, ns, not significant.

**Supplementary Figure 4:**
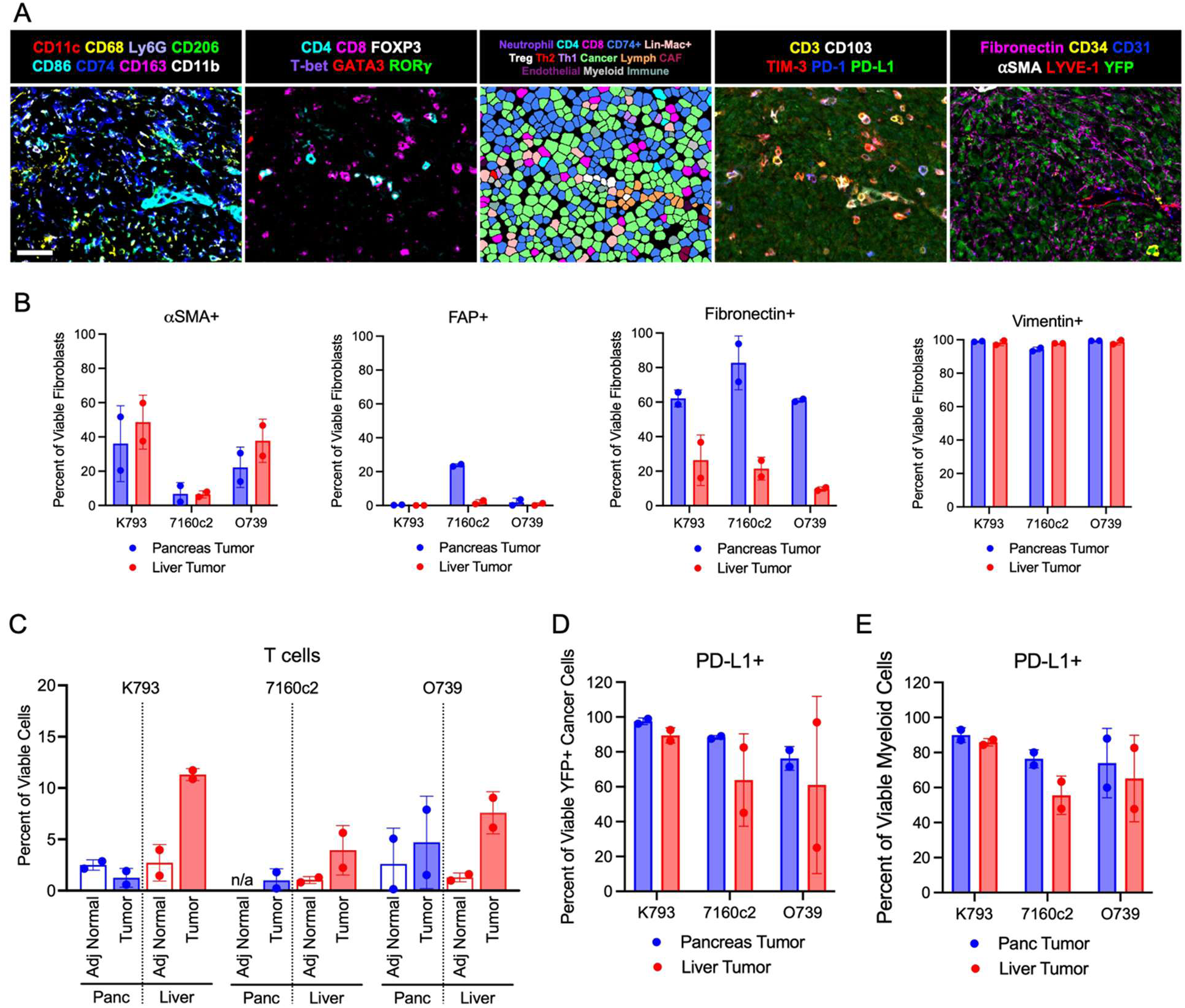
Anatomical Site Is a Major Determinant of PDAC Immune Composition. **A.** Representative micrographs of the discussed immune phenotypic markers and cancer and stroma (scale 20 µm). The middle panel displays cell phenotype overlays representing the discussed populations. **B**. Frequency of each CAF marker (αSMA, FAP, Fibronectin, Vimentin) out of viable fibroblasts according to the tumor ROI of each cell line. **C**. Frequency of T cells out of all viable cells according to ROI. **D**. PD-L1 expression of viable YFP^+^ cancer cells according to tumor ROIs. **E**. PD-L1 expression of viable myeloid cells (CD45^+^, CD19^-^, CD3^-^, NCR1^-^) according to tumor ROIs. n = 2 mice for all groups.

**Supplementary Figure 5:**
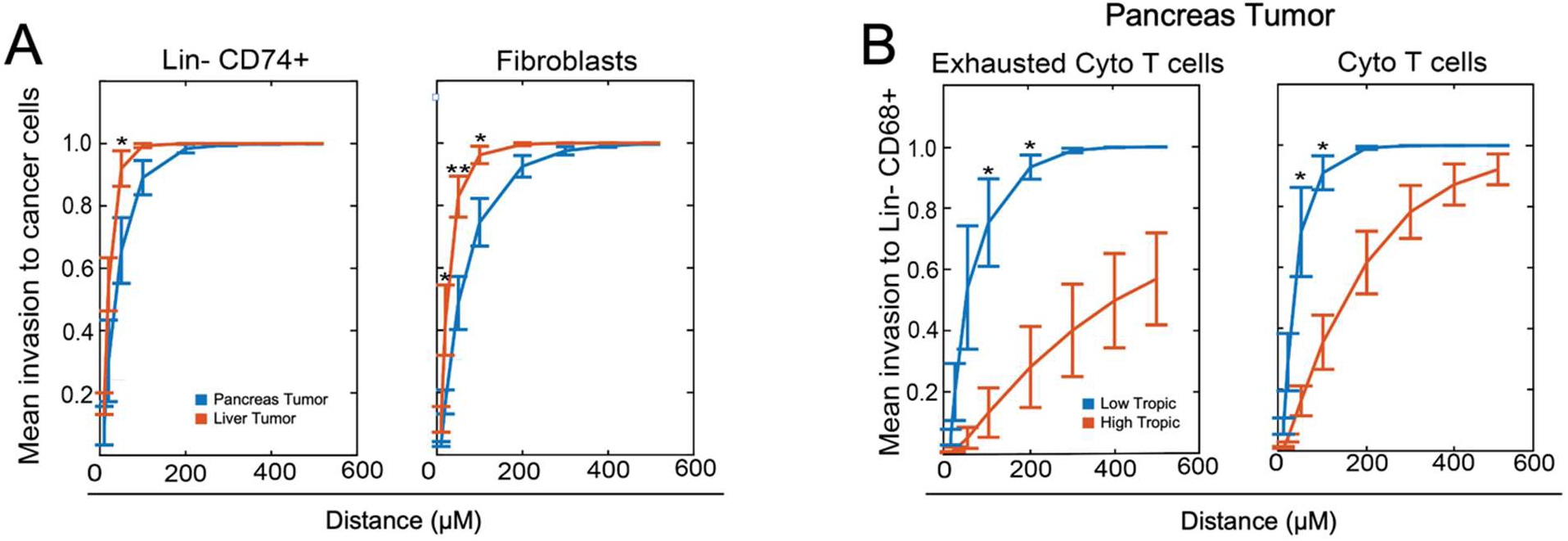
Liver tumors display enhanced TME infiltration around cancer cells. **A.** G function plots showing mean infiltration to cancer cells of Lin^-^ CD74^+^ (APC) cells and fibroblasts within pancreas and liver tumors. **B**. G function plots showing mean infiltration to Lin^-^ CD68^+^ cells from exhausted (PD-1^+^ TIM3^+^) cytotoxic T cells and total cytotoxic T cells within pancreas tumors of high liver-tropic cell lines (K793 and 7160c2) and low-liver tropic cell line (O739). Two-sample, two-sided t-tests. * p<0.05, ** p<0.01.

**Supplementary Figure 6:**
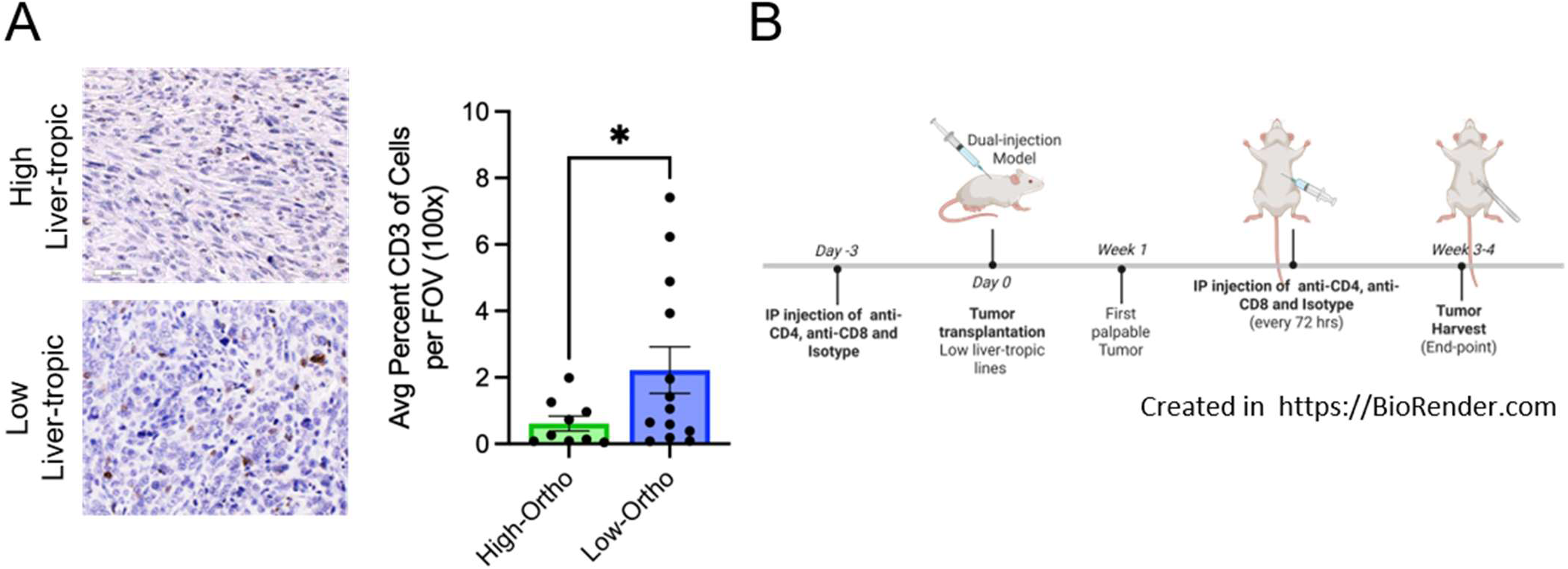
T cells constrain PDAC liver metastatic outgrowth. **A**. Representative CD3 IHC micrographs (scale 50µm) and quantification of average percent CD3^+^ cells per 100x FOV. n= High lines: 9, Low lines: 13 mice. Average of 7 FOV per mouse. Data are ± Standard Error Mean (SEM). Unpaired t test with Welch’s correction. **B**. Schematic of T cell depletion study.

